# When the ventral visual stream is not enough: A deep learning account of medial temporal lobe involvement in perception

**DOI:** 10.1101/2020.10.07.327171

**Authors:** Tyler Bonnen, Daniel L.K. Yamins, Anthony D. Wagner

## Abstract

The medial temporal lobe (MTL) supports a constellation of memory-related behaviors. Its involvement in perceptual processing, however, has been subject to enduring debate. This debate centers on perirhinal cortex (PRC), an MTL structure at the apex of the ventral visual stream (VVS). Here we leverage a deep learning framework that approximates visual behaviors supported by the VVS—i.e. lacking PRC. We first apply this approach retroactively, modeling 30 published visual discrimination experiments: Excluding non-diagnostic stimulus sets, there is a striking correspondence between VVS-modeled and PRC-lesioned behavior, while each are outperformed by PRC-intact participants. We corroborate and extend these results with a novel experiment, directly comparing PRC-intact human performance to electrophysiological recordings from the macaque VVS: PRC-intact participants outperform a linear readout of high-level visual cortex. By situating lesion, electrophysiological, and behavioral results within a shared computational framework, this work resolves decades of seemingly inconsistent findings surrounding PRC involvement in perception.

## 1 Introduction

Animal behavior is informed by previous experience. To understand how the mammalian brain supports this ability, neuroscientific data are often interpreted using two distinct cognitive constructs: ‘perception’ transforms ongoing sensory experience into behaviorally relevant abstractions (e.g. objects), while ‘memory’ enables retrieval of prior task-relevant experience. Accordingly, the ventral visual stream (VVS) is recognized for its role in visual perception (DiCarlo et al., 2012; Felleman and Van, 1991; Ullman et al., 1996), while the medial temporal lobe (MTL) is recognized for its role in memory-related behaviors (Eichenbaum and Cohen, 1993; Scoville and Milner, 1957; Squire et al., 2004). Nonetheless, an alternative taxonomy has been proposed, suggesting a more graded relationship between the behaviors supported by the VVS and MTL (Bussey et al., 2005; Murray et al., 2007). Critically, this account suggests that the MTL supports both perceptual and mnemonic behaviors (Murray and Bussey, 1999). To evaluate these competing claims, decades of experimental data have been brought to bear on whether the MTL is involved in perception (Murray and Wise, 2012; Suzuki, 2009). However, interpreting the available evidence has been subject to a longstanding debate.

This debate centers on perirhinal cortex (PRC), an MTL structure (Fig. 1a) situated at the apex of the primate VVS (Miyashita, 2019; Suzuki and Baxter, 2009). Lesion, electrophysiological, and functional imaging data have documented the role of PRC in memory-related behaviors (Aggleton and Brown, 2006; Brown et al., 2015; Miyashita, 2019; Scoville and Milner, 1957). However, early observations also revealed that PRC-related memory impairments are modulated by stimulus properties (Bussey et al., 2002; Eacott et al., 1994; Gaffan and Murray, 1992; Meunier et al., 1993), motivating perceptual experiments in PRC-lesioned primates. These early studies found that PRC-lesioned subjects were impaired on visual tasks (Buckley and Gaffan, 1997, 1998a; Buckley and Gaffan, 1998b; Bussey et al., 2002). A perceptual-mnemonic hypothesis emerged to account for these observations, suggesting that 1) PRC contributes to visual perception, independent from other regions within the MTL, and 2) subjects with PRC lesions will necessarily rely on perceptual processing in the VVS (Bussey et al., 2005). Critically, from this perspective, PRC-related perceptual impairments are only evident/expected in tasks that require sufficiently ‘complex’ visual representations (Murray and Bussey, 1999)—that is, visual representations not supported by the VVS alone.

**Figure 1:**
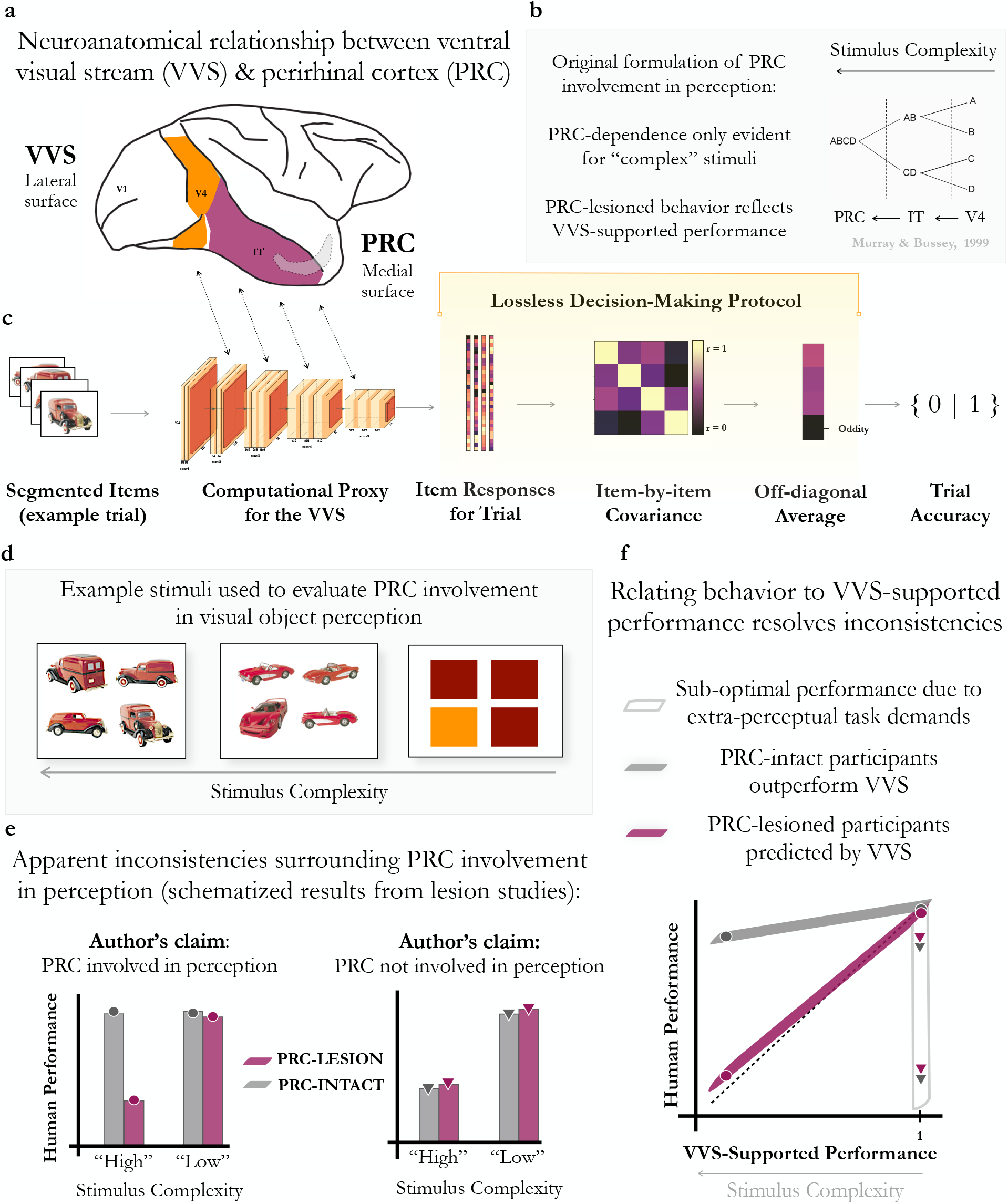
Resolving seemingly inconsistent experimental findings by situating human behavior in relationship to VVS-supported performance. **(a)** Perirhinal cortex (PRC) is a neuroanatomical structure within the medial temporal lobe (MTL) situated at the apex of the ventral visual system (VVS), downstream of ‘high-level’ visual structures such as inferior temporal (IT) cortex. **(b)** A perceptual-mnemonic hypothesis posits that PRC enables perceptual behaviors not supported by canonical sensory cortices (e.g. IT), in addition to its mnemonic functions. Critically, PRC-related perceptual impairments are only expected on so-called ‘complex’ perceptual stimuli. **(c)** Our trial-level protocol formalizes perceptual demands in ‘oddity’ visual discrimination tasks by simulating visual discrimination behaviors in the *absence* of PRC. We segment each stimulus screen containing N objects into N independent images, pass them to a computational proxy for the VVS, and extract N feature vectors from an ‘IT-like’ layer. After generating a item-by-item covariance matrix for each trial, the item with the least off-diagonal covariance is marked as the ‘oddity.’ Critically, this is a lossless decision-making protocol which is agnostic to extra-perceptual task demands (i.e. memory, attention, motivation). **(d)** Example stimuli used to evaluate the perceptual-mnemonic hypothesis that span the range of stimulus ‘complexity,’ taken from trials used in Barense et al., 2007. **(e)** Schematic of seemingly inconsistent experimental findings for (left) and against (right) PRC involvement in perception. These experiments are classified as *either* complex or control trials—a binary distinction—as is often the case in the literature. Deciding whether a stimulus set is ‘complex’ depends on experimenter discretion, not objective measures. **(f)** We propose that these apparent inconsistencies can be resolved by situating human behavior in relationship to a linear readout of the VVS: PRC-lesioned behavior is predicted by a linear readout of high-level VVS (purple) while PRC-intact behavior outperforms a linear readout of the VVS (grey). We consider experiments described as ‘complex’ but which the model performs at ceiling (i.e. x=1) to be non-diagnostic. In this formulation of the perceptual-mnemonic hypothesis, stimulus ‘complexity’ is continuous and inversely related to VVS-supported performance.

Methodological concerns were raised with the perceptual-mnemonic interpretation of PRC-lesioned behaviors. Some suggested that PRC-related deficits in these experiments were a consequence of extra-perceptual task demands (e.g. trial-level memory demands), rather than to perceptual demands (Buffalo et al., 1998a). Additionally, there were concerns that it is concurrent damage to PRC-adjacent sensory cortices, rather than to PRC, that explains perceptual deficits in some lesioned subjects (Buffalo et al., 1998a; Suzuki, 2009). Immediately adjacent to PRC is inferior temporal (IT) cortex, a region that plays a well-established role in visual object perception, lending potential credence to this concern. Moreover, analyses over a series of experiments examining the behavior of non-human primates with lesions targeting PRC revealed that visual discrimination impairments were best explained by the extent of inadvertent lesions of IT (Buffalo et al., 1998b).

To resolve these competing claims, experimentalists on both sides of the perceptual-mnemonic debate developed ‘oddity’ visual discrimination tasks; in these experiments, all requisite perceptual information is available on a single stimulus screen, in order to minimize demands on memory. In each trial, participants freely view a stimulus screen containing multiple objects (Fig. 1d), then choose the item whose identity does not match the others. Control trials (e.g. color/object category trials) are designed to rely on canonical visual perceptual cortices, irrespective of difficulty. Performance on these trials is used to indicate whether VVS structures are intact, with the goal of ruling out the possibility that perceptual impairments are due to damage in PRC-adjacent perceptual cortices. Conversely, diagnostic trials are designed to require putatively ‘complex’ perceptual representations not supported by the VVS: the presence or absence of PRC-lesioned deficits on these trials is used to evaluate PRC involvement in perception. Comparing the accuracy of PRC-intact/lesioned subjects on diagnostic trials, given the preserved performance on control trials, has been agreed upon as a suitable means to evaluate the perceptual-mnemonic hypothesis.

Unfortunately, oddity visual discrimination tasks administered to PRC-lesioned and -intact participants have generated a seemingly inconsistent pattern of experimental evidence. More pointedly, results from these studies have been used both to support (Barense et al., 2007; Buckley et al., 2001; Bussey et al., 2003; Inhoff et al., 2019; Lee et al., 2006; Lee et al., 2005b) and refute (Knutson et al., 2012; Levy et al., 2005; Squire et al., 2006; Stark and Squire, 2000) the perceptual-mnemonic hypothesis (inconsistent outcomes schematized in Fig. 1e). In response, both sides of the debate have argued that there are oddity visual discrimination experiments that have been mistakenly used to evaluate the role of PRC in perception. Critics of the perceptual-mnemonic hypothesis have continued to claim that PRC-related deficits on oddity tasks are due to memory-related task demands and possible damage to PRC-adjacent visual cortex, while proponents have argued that the absence of PRC-related deficits are due to ‘insufficiently complex’ stimulus sets. However, it has not been possible to arbitrate between these competing claims, given the lack of formal methods for determining perceptual and mnemonic task demands.

We suggest that these apparent inconsistencies can be resolved by situating behavior in relation to perceptual processing supported by the VVS. Experimental accuracy supported by a linear readout of the VVS offers a direct assessment of perceptual processing in the absence of extra-perceptual task demands; we refer to this lossless readout as ‘VVS-supported performance.’ In this framework, stimulus ‘complexity’ is continuous and inversely related to VVS-supported performance (Fig. 1f). A perceptual-mnemonic hypothesis would predict that experimental observations within the literature fall into three distinct distributions along this axis. First, PRC-lesioned behavior is approximated by VVS-supported performance (Fig. 1f: purple). Second, PRC-intact participants outperform the VVS (Fig. 1f: grey). And third, experiments where VVS-supported performance is at ceiling and no additional (e.g. PRC-dependent) perceptual processing should be necessary (Fig. 1f: white). By design, control trials belong to this third distribution, and are accepted as non-diagnostic, where any below-ceiling performance is attributed to extra-perceptual task demands. By the same logic, regardless of whether a stimulus has been *described* as ‘complex,’ it belongs in this third distribution if VVS-supported performance is at ceiling. Thus, situating human behavior in relation to VVS-supported performance provides an account of when perception depends on processes beyond the VVS (e.g. PRC-dependent) and, conversely, identifies when performance impairments may be due to extra-perceptual task demands.

We evaluate this formulation of the perceptual-mnemonic hypothesis by situating lesion, electrophysiological, and behavioral results within a shared computational framework. First, we analyze a ‘retrospective’ dataset composed of 30 published oddity experiments administered to PRC-lesioned and -intact human participants. As neural recordings from the VVS are not available for these experiments, we leverage a model class that is able to predict neural activity throughout the VVS: task-optimized convolutional neural networks (Bashivan et al., 2019; Yamins et al., 2014). We develop an analytic approach that generates trial-by-trial predictions of VVS-supported performance on oddity tasks, directly from experimental stimuli (Fig. 1c). This ‘stimulus-computable’ framework enables us to approximate behaviors supported by the VVS, without reliance on informal, descriptive accounts of experimental stimuli. We deploy this computational approach to estimate VVS-supported performance on each experiment within this retrospective dataset, evaluating the perceptual-mnemonic hypothesis through a series of increasingly stringent analyses. Next, we develop and assess a novel, model-driven experiment that enables us to evaluate the perceptual-mnemonic hypothesis with unprecedented resolution: we compare human behavior on oddity visual discrimination tasks directly to electrophysiological recordings from high-level visual cortex in the macaque. Collectively, this work situates evidence from multiple experimental settings within a unified computational framework, resolving decades of *seemingly* inconsistent experimental outcomes surrounding PRC involvement in visual object perception.

## 2 Results

### 2.1 Retrospective Analysis

Through a comprehensive literature review we identify oddity visual discrimination studies administered to PRC-intact and -lesioned human participants (STAR Methods: Literature Review). We assemble a ‘retrospective dataset’ composed of stimuli and behavioral data from 30 published experiments, including studies used as evidence for and against the perceptual-mnemonic hypothesis (STAR Methods: Retrospective Dataset). With a task-optimized convolutional neural network, we identify a model layer that best fits electrophysiological responses collected from IT-cortex in the macaque (STAR Methods: Model Fit to Electrophysiological Data). Using an unweighted, linear decoder off model responses from this ‘IT-like’ layer to solve each trial, we then average accuracy across trials in each experiment (STAR Methods: Model Performance on Retrospective Dataset). Thus, for each experiment in the retrospective dataset, we have a single value corresponding to the averaged performance that would be expected by a linear readout of high-level visual cortex— absent extra-perceptual task demands. We refer to this estimate as ‘model performance.’ As a final analysis of the retrospective dataset, we estimate what stage of VVS processing PRC-lesioned performance is reliant on. We note that all key findings reported in the retrospective dataset are consistent across all model instances evaluated; we report results from a single model for simplicity, and refer to Supplemental Figures S3 and S7 for results across model instances.

#### 2.1.1 Model performance: A computational proxy for high-level visual cortex

We begin with a task-optimized convolutional neural network, pre-trained to perform object classification. Using previously collected electrophysiological responses from macaque VVS (Majaj et al., 2015), we identify a model layer that best fits high-level visual cortex: Given a set of images, we learn a linear mapping between model responses and a single electrode’s responses, then evaluate this mapping using independent data (STAR Methods: Model Fit to Electrophysiological Data). For each model layer, this analysis yields a median cross-validated fit to noise-corrected neural responses, for both V4 and IT—corresponding to earlier and later stages of processing within the VVS. As is consistently reported (Rajalingham et al., 2018; Schrimpf et al., 2020; Yamins et al., 2014), early model layers (i.e. first half of layers) better predict neural responses in V4 than do later layers (unpaired ttest: *t*(8) = 2.70, *P* = .015), while later layers better predict neural responses in IT, a higher-level region (unpaired ttest: *t*(8) = 3.70, *P* = .002). Accordingly, differential fit to IT cortex (Δ_*IT-V*4_) increases with model depth (ols regression: *β* = .98, *F* (1, 17) = 20.91, *P* = 10^*-*13^). Peak V4 fits occur in model layer pool3 (noise-corrected *r* = .95 ± .30STD) while peak IT fits occur in con5 1 (noise-corrected *r* = .88 ± .16STD). Having identified this ‘IT-like’ layer, we use an unweighted, linear decoder off this model layer to solve each trial in our initial analyses; for comparison with human behavior, we average performance across trials on each experiment (STAR Methods: Model Performance on Retrospective Dataset). Here we use the term ‘experiment’ in a way that is interchangeable with ‘condition’ not ‘study.’ For example, a single study typically contains multiple experiments.

#### 2.1.2 Identification of non-diagnostic stimulus sets

Our approach identifies 14 experiments that were used to evaluate PRC involvement in perception, yet model performance is at ceiling (Fig. 2 at x=1, corresponding to 100% accuracy). We suggest these experiments are not diagnostic of PRC involvement in perception: Oddity experiments perfectly supported by the VVS are not positioned to evaluate the role of PRC in perception, as no additional *perceptual* processing should be required (See STAR Methods: Non-Diagnostic Experiments for more details about this reasoning). These ‘non-diagnostic’ stimuli include eight experiments in which performance did not differ between PRC-intact and -lesioned participants (1 experiment in Buffalo et al., 1998a, all 7 experiments in Knutson et al., 2012; Supplemental Fig. S1a-b). The original authors suggest these experiments provide evidence against PRC involvement in perception. In another six experiments, performance differed between PRC-lesioned and -intact subjects (all 3 ‘Fribble’ experiments in Barense et al., 2007, all 3 ‘Face Morphs’ in Inhoff et al., 2019; Supplemental Fig. S1c-d). The original authors suggest these experiments provide evidence in support of PRC involvement in perception. While our modeling approach is not designed to provide an account of performance on these non-diagnostic experiments, we speculate as to the divergent outcomes across experiments in the Supplement.

**Figure 2:**
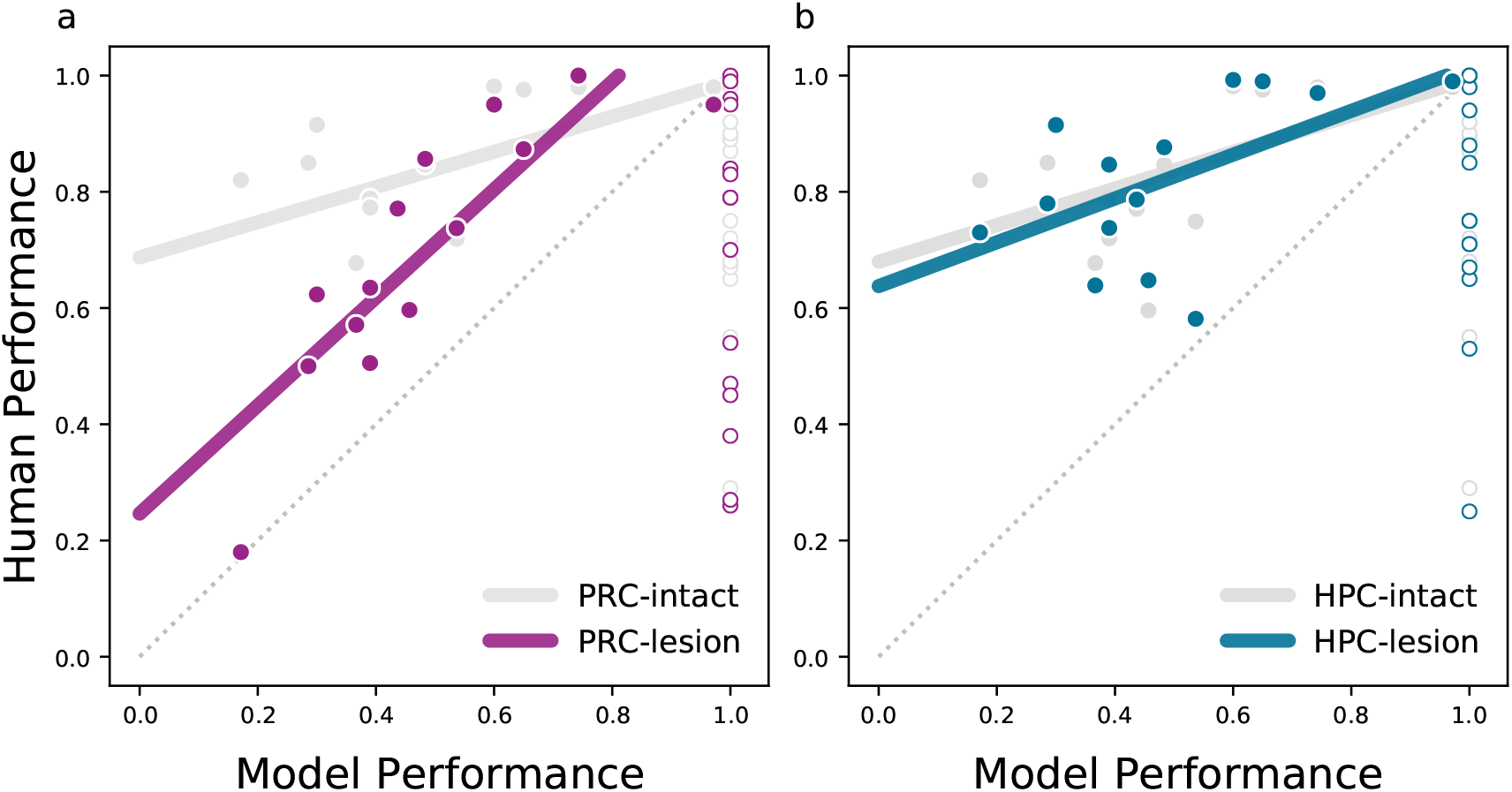
A computational proxy of the VVS predicts PRC-lesioned performance, while both are outperformed by PRC-intact participants. We collect previously published ‘oddity’ experiments administered to PRC-lesioned and -intact human participants. We present experimental stimuli to a computational proxy for the VVS, then use a linear decoder off an ‘IT-like’ layer to predict the oddity in each trial. We then average performance across all trials in each experiment: This single value (‘model performance’) corresponds to the experimental accuracy expected from a linear readout of IT cortex under a lossless decision-making protocol. Stimuli where model performance is at ceiling (x=1, open dots) are not relevant for evaluating the role of PRC in perception: As VVS responses should support perfect discrimination between stimuli, any below-ceiling performance in the human is attributed to extra-perceptual task demands (e.g. memory). **(a)** This computational proxy for IT cortex predicts the behavior of PRC-lesioned participants, while PRC-intact participants outperform both model and PRC-lesioned participants. **(b)** HPC-lesioned and intact participants all outperform this computational model on relevant stimuli; both for participants with an entirely intact medial temporal lobe, which includes PRC, as well as participants with selective damage to the hippocampus that spare PRC. Together, these results suggest that PRC-lesioned behavior reflects a linear readout of the VVS, neurotypical behaviors on these tasks outperform the VVS, and this behavior is dependent on PRC.

We note that these results are not dependent on the particular model we use to evaluate these stimuli. All computational proxies for the VVS identify the same experiments as non-diagnostic (STAR Methods: Non-Diagnostic Experiments). Additionally, pixel-level analyses identify the same experiments as non-diagnostic: If, instead of model responses, we run an identical analysis over the original images flattened into vector form, we achieve 100% accuracy on all model-identified non-diagnostic stimuli, while performing below 100% on all model-identified diagnostic experiments. That is, non-diagnostic experiments can be performed by evaluating pixel-level differences between stimuli, whereas diagnostic experiments cannot be solved in this manner. Finally, we note that all non-diagnostic experiments used arrays of stimuli from the same viewpoint (see Supplemental Fig. S2), making them visually distinct from those experiments where model performance was below ceiling (see Supplemental Fig. S4). Taken together, these findings suggest that experiments exist in the literature that may not be diagnostic of PRC involvement in perception. While this concern has been consistently raised in the literature, our work offers the first stimulus-computable operationalization of these predictions. We exclude these experiments from further analyses.

#### 2.1.3 PRC-lesioned subjects are impaired on oddity visual discrimination tasks

To make claims about PRC involvement in oddity visual discrimination behaviors, we are principally interested in the comparison between PRC-lesioned behavior and their non-lesioned, age and IQ matched controls (i.e. ‘PRC-intact’). However, human PRC lesions are often accompanied by damage to other prominent structures within the MTL, such as the hippocampus (HPC). To ensure that behavioral impairments are a consequence of damage to PRC and not HPC, we also compare the behavior of participants with selective hippocampal damage (i.e. ‘HPC-lesioned’) to their non-lesioned, age and IQ matched controls (i.e. ‘HPC-intact’)—where both HPC-lesioned and HPC-intact participants have an intact PRC. This is standard practice in the MTL literature. Across the 14 diagnostic experiments in the retrospective dataset PRC-lesioned participants are significantly impaired relative to PRC-intact participants (paired ttest, *β* = .14, *t*(13) = 2.68, *P* = .019), while HPC-lesioned participants show no such impairment (paired ttest, *β* = .01, *t*(13) = .73, *P* = .479). Directly comparing the difference between PRC-intact/lesioned participants with HPC-intact/lesioned participants, there is a significant difference between lesioned groups (*β* = .13, *F* (1, 26) = 2.34, *P* = .028). PRC-intact participants perform significantly better on oddity experiments than PRC-lesioned participants, while there is no such difference between HPC-intact and -lesioned participants.

#### 2.1.4 A computational model of the VVS approximates PRC-lesioned performance

The previous section demonstrates a coarse distinction between PRC-lesioned and -intact performance. A more incisive test of the perceptual-mnemonic hypothesis would be to predict the relative impairments observed between PRC-lesioned and -intact participants. To this end, we compare model and human performance across diagnostic experiments in the retrospective dataset (STAR Methods: Model Performance on Retrospective Dataset). First, we observe a clear correspondence between PRC-lesioned behavior and model performance; as our estimate of VVS-supported performance decreases, so too does PRC-lesioned accuracy (Fig. 2a, purple; ols regression *β* = .85, *F* (1, 12) = 5.59, *P* = 1×10^*-*4^). Next, we find that there is not a statistically significant relationship between PRC-intact accuracy and a computational proxy for the VVS (Fig. 2a, grey: ols regression *β* = .85, *F* (1, 12) = 2.05, *P* = .063); these participants outperform the model (ols regression *β* = .35, *t*(13) = 7.32, *P* = 6×10^*-*6^). Critically, when predicting human accuracy from model performance, we observe a significant interaction between PRC-intact and PRC-lesion groups (ols regression *β* = .63, *F* (3, 24) = 2.82, *P* = .010). This interaction is not observed for the hippocampal groups (HPC-lesion/HPC-intact *β* = .06, *F* (3, 24) = .28, *P* = .781). To be more explicit about the interpretation of this differential correspondence between model performance and PRC-lesioned behavior, we take the difference between PRC-intact and -lesioned participants, resulting in a difference score for each experiment. This difference is predicted by model performance (ols regression *β* = -.63, *F* (1, 12) = -3.17, *P* = .008) with the sign indicating that as model performance is degraded, the difference between PRC-intact and -lesioned participants increases.

Collectively, our results demonstrate that computational proxies of the VVS predict PRC-related deficits observed across experiments, confirming a key prediction of the perceptual-mnemonic hypothesis. We note that these findings are consistent across model instances: All state-of-the-art models evaluated demonstrate the same differential fit to PRC-lesioned participants illustrated by the interaction analysis above (Supplemental Fig. S3). Nonetheless, we find a mean difference between model and PRC-lesioned performance, with lesioned participants outperforming the model (paired ttest; *β* = .21, *t*(13) = 6.63, *P* = 2×10^*-*5^). Three sources of variance may contribute to this difference: The extent of damage to PRC in these naturally occurring lesions, the model instance used to approximate VVS-supported performance (as evident in Supplemental Fig. S3), and the readout used to estimate model performance (STAR Methods: Model Performance on Novel Stimuli & Comparing Readout Performance). We elaborate on these sources of variance in the Methods and Supplement.

#### 2.1.5 PRC-lesioned oddity performance appears to rely on high-level visual cortex

The perceptual-mnemonic hypothesis claims that PRC-lesioned behavior relies on the VVS (Murray and Bussey, 1999). Here we seek to determine which structures within the VVS PRC-lesioned behavior is reliant on. Historically, this question has been raised through concerns that PRC-lesioned deficits on oddity tasks are a consequence of concurrent damage to PRC-adjacent perceptual regions, such as IT cortex (Buffalo et al., 1999). Within our retrospective analysis, we find that IT-like model layers exhibit the same relative deficits as PRC-lesioned subjects, suggesting that IT-like representations do not enable PRC-intact levels of performance. However, to claim that PRC-lesioned participants rely on IT cortex requires additional evidence: High-level visual cortex must predict PRC-lesioned behavior significantly better than earlier stages of visual processing. Otherwise it is not possible to isolate early-from late-stage VVS dependence. Thus, as a final evaluation of the retrospective dataset, we determine whether high-level visual cortex uniquely explains PRC-lesioned performance. Lacking direct electrophysiological recordings from early- and late-stage processing from within the VVS, we again leverage a computational proxy for the primate VVS. After identifying model layers that best fit V4 and IT cortex (as per the model evaluation methods outlined in Model performance: A computational proxy for high-level visual cortex) we generate a metric to describe each layer’s differential fit to macaque IT cortex (IT fit – V4 fit: Δ_*IT-V*4_, Fig. 3a). Just as in previous analyses, we determine model performance on each (diagnostic, n=14) experiment within the retrospective dataset, but now perform this analysis using each model layer, not only an ‘IT-like’ layer (several layers visualized in Fig. 3b: PRC-lesion/intact top, HPC-lesion/intact bottom). With these model performance-by-layer estimates we generate a metric for the relative fit to PRC-lesioned (Δ_*prc*_ = *MSPE*_*prc*.*intact*_ - *MSPE*_*prc*.*lesion*_, STAR Methods: VVS Reliance) and HPC-lesioned (Δ_*hpc*_) performance for each model layer. We relate these metrics from the retrospective dataset (Δ_*prc*_ and Δ_*hpc*_) to the model’s differential fit to macaque IT cortex (Δ_*IT-V*4_).

**Figure 3:**
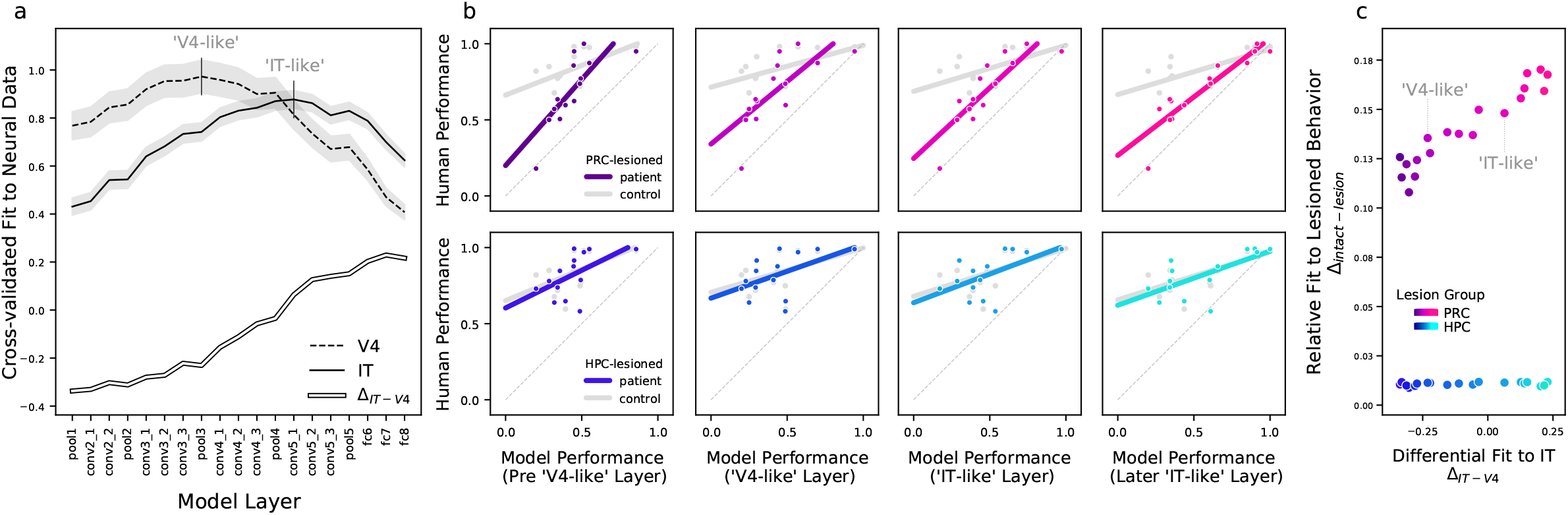
PRC-lesioned oddity performance appears to rely on high-level visual cortex. The perceptual-mnemonic hypothesis claims that PRC-lesioned behavior relies on the VVS (Murray and Bussey, 1999), and some evidence suggests that PRC-lesioned subjects rely on inferior temporal (IT) cortex (Buffalo et al., 1998a). To evaluate this possibility, we leverage our computational proxy for the VVS to estimate the relative contributions that V4- and IT-supported behaviors have on the retrospective dataset: **(a)** Using the model’s cross-validated fit to electrophysiological recordings, we determine each layer’s fit to IT (solid) and V4 (dashed) cortex. This enables us to identify the most ‘IT-like’ and ‘V4-like’ layer within the model (vertical lines, labeled). We also determine the degree to which each layer better predicts IT, relative to V4, by estimating the differential fit to IT cortex– computing the difference between IT and V4 neural fits (Δ_*IT-V*4_: hollow). **(b)** To evaluate whether IT-like layers uniquely explain PRC-lesioned behaviors, we determine model performance on the retrospective dataset, across all layers, and evaluate each layer’s fit to human performance. Model performance from all layers predicts PRC-lesioned performance, including performance from V4- and IT-like layers. Additionally, (as in previous analysis, Fig. 2) we observe a significant interaction between PRC-lesioned and -intact subjects when predicted by most model layers (top). There is not an interaction between HPC-lesioned and -intact subjects for any model layers (bottom). These results suggest that both V4-like (2nd column) and IT-like (3rd column) model layers can serve as a computational proxy for PRC-lesioned performance. **(c)** Nonetheless, we evaluate whether IT-like layers increase their fit to PRC-lesioned behaviors: We compare each layer’s differential fit to IT cortex (Δ_*IT-V*4_, x axis) to that layer’s relative fit to lesion behavior (Δ_*lesion*_, y axis). For each layer, we determine the model’s relative fit to lesioned behavior (Δ_*lesion*_), using the mean squared prediction error (MSPE): We first compute the MSPE between the model and each group, then determine the difference between lesioned and intact participants, for both PRC and HPC groups (Δ_*prc*_ and Δ_*hpc*_, respectively). While all model layers predict PRC-lesioned subject performance (as evident in b), model layers that better fit IT cortex exhibit quantitatively (but not qualitatively) better fits to PRC-relevant behavior (Δ_*prc*_, top). There is no relationship with HPC-lesioned behavior (Δ_*hpc*_, bottom). These results are suggestive that PRC-lesioned performance reflects a linear readout of high-level visual cortex, but are inconclusive, as there is not a clear separation between model performance from IT-like and V4-like layers. These data highlight a limitation of the stimuli used to evaluate PRC-lesioned behavior in the retrospective dataset.

Our model performance-by-layer analyses reveal two outcomes. First, model performance from all layers predicts PRC-lesioned performance, including V4- and IT-like layers (i.e. pool3, *β* = .74, *F* (1, 12) = 3.86, *P* = .002, and conv 51, *β* = .85, *F* (1, 12) = 5.59, *P* = 10^*-*4^, respectively, as can be seen in Fig. 3b, top). Moreover, the interaction between PRC-lesioned and -intact performance predicted by the perceptual-mnemonic hypothesis (i.e. Fig. 2a) is observed across most model layers, including V4- (*β* = .55, *F* (3, 24) = 2.08, *P* = .048) and IT-like layers (*β* = .63, *F* (3, 24) = 2.82, *P* = .010). Conversely, an interaction between HPC-lesion and -intact performance was not observed at any model layer (Fig. 3b, bottom). These results suggest that, for stimuli in the retrospective dataset, PRC-lesioned performance might be similar if participants were relying on V4-like (Fig. 3b, 2nd column, top) or IT-like (Fig. 3b, 3rd column, top) representations. Second, layers that better fit IT cortex are nonetheless better quantitative fits to PRC-lesioned performance: differential correspondence with macaque IT predicts relative fit to PRC-lesioned human behavior (Fig. 3c, top: ols regression *β* = .95, *F* (1, 17) = 13.20, *P* = 2 × 10^*-*10^). Across experiments, this result is especially prominent for ‘less complex’ experimental stimuli (Supplemental Fig. S6).

Moreover, when evaluating the interaction between PRC-lesion/intact behavior, in relation to model performance, only IT-like model layers are significant after correcting for multiple comparisons. Nonetheless, IT-like model layers are not significantly better at predicting PRC-lesioned participant behavior than V4-like model layers (*β* = -.11, *F* (3, 24) = -.40, *P* = .693). These data suggest that PRC-lesioned behavior may be better explained by high-level visual cortex. However, lacking a significant difference between V4-like and IT-like model performance across experiments, the retrospective dataset does not enable incisive claims about VVS-reliance in a PRC-lesioned state.

#### 2.1.6 Retrospective summary & limitations

Results from the retrospective dataset suggest that PRC-lesioned performance reflects a linear readout of the VVS. In contrast, PRC-intact behaviors outperform both PRC-lesioned participants and a computational proxy for the VVS. While this analysis resolves fundamental questions at the center of the perceptual-mnemonic debate, there are numerous limitations. First, our analysis and conclusions depend on a computational proxy for the VVS, not direct neural recordings. Second, the available stimulus sets sparsely sample the range of VVS-supported behaviors; this is due, in part, to low experimental *N* s (both experiments and participants), and reliance on experimental averages when fitting to human behavior (i.e. comparing mean performance across experiments, not trial-by-trial behaviors). Finally, there is a considerable amount of hypothesis-orthogonal variability across studies; for example, the number of stimuli used on each trial varies from 3 to 9 objects across experiments in the retrospective dataset. We develop a novel, model-based experiment that enables us to circumvent these limitations.

### 2.2 Novel Dataset

Here we compare PRC-intact human behavior *directly* to VVS-supported performance, situating human behavior in relation to electrophysiological recordings collected from high-level visual cortex in the macaque. This enables us to evaluate a central tenet of the perceptual-mnemonic hypothesis with unprecedented resolution, as well as the opportunity to address limitations in the retrospective analysis: we design an experiment that enables item-level performance estimates, continuously samples the space of stimulus ‘complexity,’ and clearly disentangles multiple stages of processing (i.e. IT vs. V4) across the VVS from PRC-intact behavior. These experiments minimize off-hypothesis experimental variance, using the minimum object configuration in each trial (*N* = 3).

#### 2.2.1 High-throughput human psychophysics experiments

We begin with stimuli that have been previously shown to separate V4-from IT-supported behavior (Majaj et al., 2015), reconfiguring these images into 3-way, within-category, oddity trials (STAR Methods: Novel Stimulus Set Generation; for examples see Fig. 4a). We develop a novel estimate of ‘model performance’ on these oddity tasks: a weighted, linear readout from an ‘IT-like’ model layer, learned via a leave-one-out cross-validated protocol (STAR Methods: Model Performance on Novel Stimuli). We administer these stimuli to PRC-intact human participants (*N* = 297) via high-throughput psychophysics experiments (STAR Methods: High-throughput Psychophysics Experiments). Finally, using the approach developed to estimate a weighted model performance, we determine the performance on these oddity trials that would be supported from a weighted readout of macaque IT and V4 (protocol illustrated in Fig. 4b). Thus, for the same stimuli, we are able to compare model performance and PRC-intact human behavior, alongside the accuracy supported by a weighted, linear readout of electrophysiological responses collected from macaque IT and V4 (Fig. 4c).

**Figure 4:**
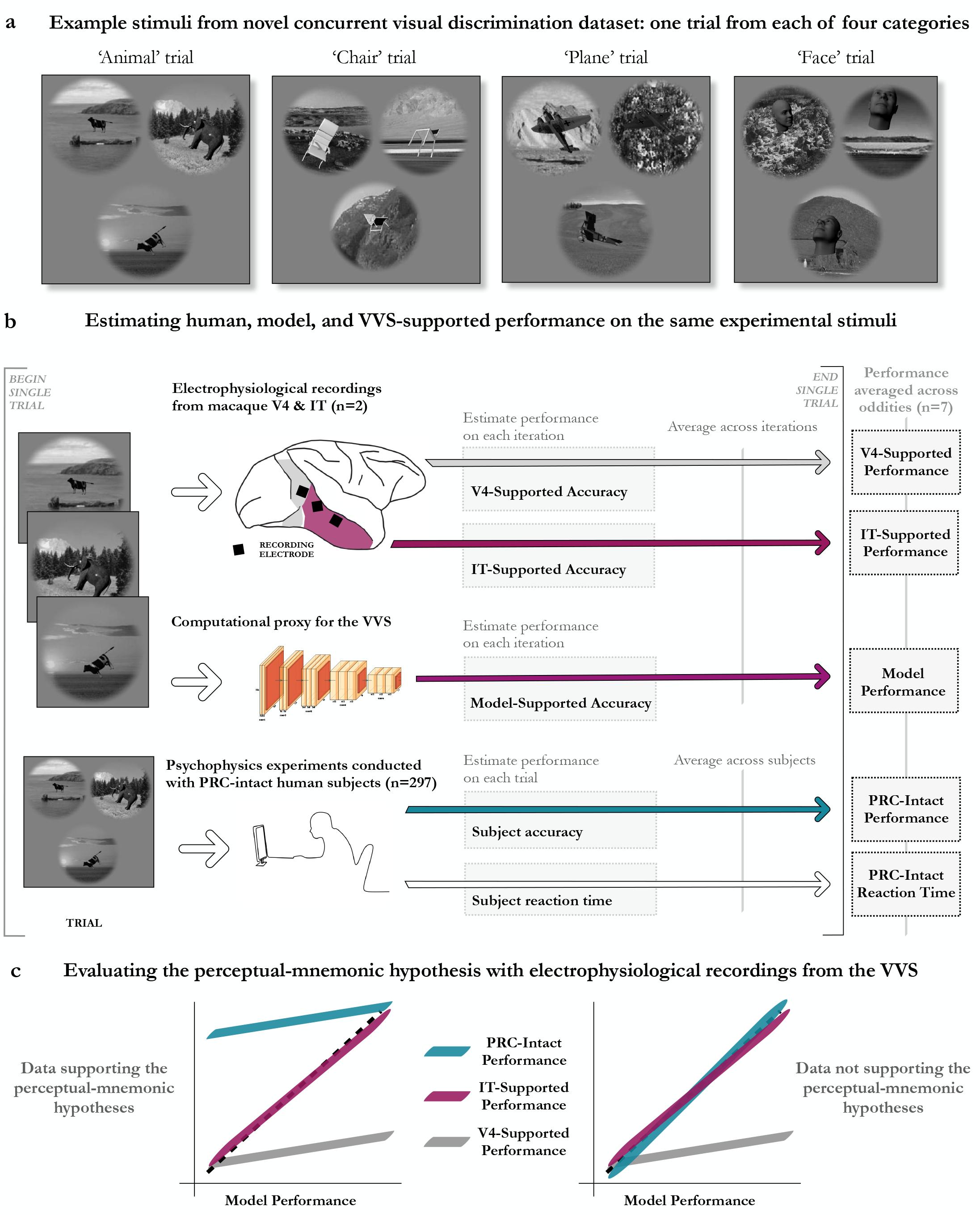
Evaluating the perceptual-mnemonic hypothesis with electrophysiological recordings from the VVS. To address limitations in the retrospective analysis, we design a novel experiment that enables item-level performance estimates, continuously samples the space of stimulus ‘complexity,’ and clearly disentangles multiple stages of processing (i.e. IT vs. V4) across the VVS. These experiments minimize off-hypothesis experimental variance, using the minimum configuration of objects in each trial (*N* = 3). Critically, this novel experiment enables us to compare PRC-intact human behavior directly to the performance supported by electrophysiological recordings from high-level visual cortex. **(a)** Example trials from the four categories in this novel oddity experiment. We generate stimuli using a computational proxy for the VVS, selecting trials that uniformly sample the space of VVS-supported performance (i.e. chance to ceiling model performance). **(b)** Our protocol enables us to evaluate PRC-intact human behavior alongside electrophysiological recordings collected from the macaque. To estimate V4- and IT-supported behavior (top), we use a modified leave-one-out cross-validated approach, averaged across multiple iterations. We use the same protocol to estimate model performance (middle). Finally, we collect human accuracy and reaction time data in a pool of online subjects (n=297; bottom). We report the item-level estimates (i.e. averaging across all oddities, for a given item) across all measures. **(c)** We use model performance to situate the electrophysiological and behavioral data within a common framework/axis: Evidence consistent with the perceptual-mnemonic hypothesis (left) predicts PRC-intact subjects will outperform high-level visual cortex, while a strictly mnemonic interpretation of PRC function (right) predicts no divergence between PRC-intact/lesioned behavior.

#### 2.2.2 PRC-intact participants outperform electrophysiological recordings from IT

PRC-intact human behavior outperforms a linear readout of macaque IT on this novel stimulus set (Fig 5c: paired ttest *β* = .24, *t*(31) = 9.50, *P* = 1×10^*-*10^), while IT significantly outper-forms V4 (Fig. 5a: paired ttest *β* = .18, *t*(31) = 6.56, *P* = 2×10^*-*7^). A computational proxy for IT demonstrates the same pattern as electrophysiological recordings, predicting IT-supported performance (Fig 5d, purple: ols regression *β* = .81, *F* (1, 30) = 13.33, *P* = 4×10^*-*14^), outper-forming V4 (Fig 5d, grey: paired ttest *β* = .26, paired ttest *t*(31) = 8.02, *P* = 5×10^*-*9^), and being outperformed by PRC-intact participants (Fig 5d, teal: paired ttest *β* = .16, *t*(31) = 5.38, *P* = 7×10^*-*6^). Nonetheless, we expect that IT-supported performance will relate to PRC-intact accuracy: For trials where IT-supported performance is greater, there is less of a demand for extra-IT processing, and more accurate PRC-intact performance. In line with this expectation, there is a reliable relationship between IT-supported and PRC-intact performance (ols regression *β* = .26, *F* (1, 30) = 8.49, *P* = 2×10^*-*9^).

**Figure 5:**
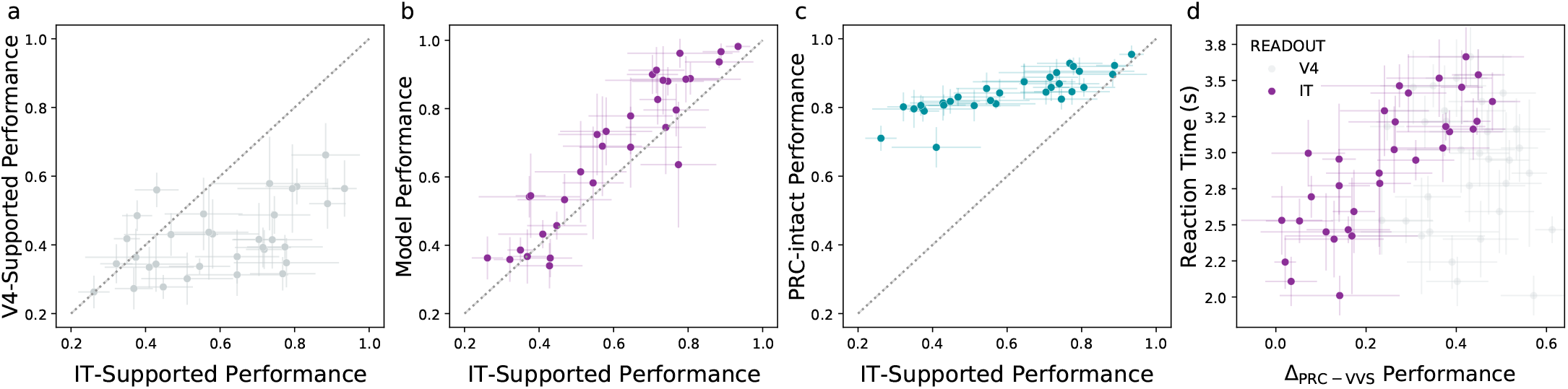
PRC-intact participants outperform a direct readout of high-level visual cortex. We directly compare PRC-intact human behaviors to electrophysiological recordings from the primate VVS using a novel stimulus set. This enables us to determine whether PRC-intact behaviors are able to outperform a *direct* readout of high-level visual cortex, while also addressing limitations within the retrospective dataset. In all plots, error bars indicate the standard deviation from the mean. **(a)** Unlike stimulus sets used within the retrospective analysis (Fig. 3b), here we can clearly separate IT-from V4-supported performance: a linear readout of IT cortex (i.e. IT-supported performance) outperforms a weighted, linear readout of V4 (grey points below the diagonal). **(b)** Our computational proxy for IT cortex is able to predict IT-supported performance, further validating the model approach used in previous analyses. **(c)** Critically, PRC-intact human participants outperform IT-supported behaviors. This is the first direct comparison of PRC-intact performance on oddity tasks in relation to electrophysiological recordings from high-level visual cortex, confirming a central tenet of the perceptual-mnemonic hypothesis. **(d)** Interestingly, we find that the difference between PRC-intact and IT-supported performance scales linearly with reaction time; on each item, subjects require more time in order to outperform a linear readout of IT. These data confirm predictions central to the perceptual-mnemonic hypothesis with unprecedented resolution, validate the computational results on the retrospective dataset, and characterize the temporal dynamics of putatively PRC-dependent visual behaviors.

In addition to these accuracy metrics, we find that the PRC-intact human reaction time for each item is a reliable predictor of IT-supported performance: longer reaction times are observed for items with lower IT-supported accuracy (ols regression *β* = -.88, *t*(31) = -10.00, *P* = 4×10^*-*11^). Framed more explicitly, for each item, the difference between IT-supported and PRC-intact per-formance is predicted by reaction time (Fig. 5d, purple: ols regression *β* = .81, *F* (1, 31) = 7.44, *P* = 3×10^*-*8^). This relationship is also observed for model performance (ols regression *β* = .72, *F* (1, 31) = 5.62, *P* = 4×10^*-*6^) but not V4-supported performance (Fig. 5d, grey: ols regression *β* = -.08, *F* (1, 31) = -.41, *P* = .682) such that there is a significant interaction between regions (i.e. PRC-intact – IT-supported and PRC-intact – V4-supported) when predicting reaction time (ols regression *β* = .49, *F* (3, 60) = 3.41, *P* = .001). These results demonstrate that PRC-intact human participants require more time to choose among items that are not linearly separable in IT, in a way that scales inversely with IT-supported performance.

## 3 Discussion

For decades, experimentalists have sought to determine the involvement of PRC in visual perception, resulting in a pattern of seemingly inconsistent experimental outcomes (Buffalo et al., 1998a; Bussey et al., 2005; Murray and Bussey, 1999; Suzuki, 2009). Using a computational proxy for the primate visual system, validated with electrophysiological recordings from the macaque, we developed a computational approach that formalizes longstanding claims within this literature. We first deploy this approach on a ‘retrospective dataset’ of 30 published oddity experiments administered to PRC-lesioned and -intact human participants. This dataset includes seminal studies used to argue both sides of the perceptual-mnemonic debate, requiring that our computational approach account for these seemingly conflicting findings.

First, we report that numerous experiments within the retrospective dataset may not be diagnostic of PRC involvement in perception. We identify and exclude these non-diagnostic experiments by operationalizing longstanding claims within the perceptual-mnemonic literature (Barense et al., 2007; Buckley et al., 2001; Inhoff et al., 2019; Lee et al., 2006; Lee et al., 2005b; Levy et al., 2005; Murray and Baxter, 2006; Murray and Bussey, 1999): If a computational proxy for the VVS is able to perform these experiments with 100% accuracy, this suggests that no perceptual processing beyond the VVS is necessary. Not only do all models evaluated identify the same experiments as non-diagnostic, but a pixel-level analysis (in place of model responses) also performs these experiments—and only these experiments—with 100% accuracy (STAR Methods: Non-Diagnostic Experiments). Perhaps surprisingly, these experiments are found on both sides of the debate (Supplemental Fig. S1), suggesting that non-diagnostic stimuli may have masked the presence (and absence) of PRC-related deficits in visual object perception. While our modeling approach is not designed to account for the relative patterns of performance on non-diagnostic tasks, we provide a speculative account of these patterns in the Supplement.

Next, we evaluate the behavior of PRC-lesioned and -intact performance on diagnostic experiments. Directly comparing subject groups, we find that PRC-lesioned participants are significantly impaired relative to their PRC-intact counterparts. Moreover, we observe a striking correspondence between computational proxies for the VVS and PRC-lesioned human behavior. Conversely, PRC-intact participants substantially outperform PRC-lesioned participants and model performance. In a critical test of the perceptual-mnemonic hypothesis, we observe a significant between-group interaction: PRC-lesioned performance is significantly better fit by computational proxies for the VVS than is PRC-intact performance (Fig. 2). These results confirm key predictions of the perceptual-mnemonic hypothesis: PRC appears to provide an advantage over the VVS in performance on oddity visual discrimination tasks. Moreover, this work seamlessly accounts for the variance in PRC-lesioned deficits observed across experiments in the available literature: PRC-lesioned deficits fall along a continuum, tracking inversely with VVS-supported performance, suggesting that the available experiments place differential demands on PRC. Meanwhile, there are no such impairments observed for hippocampal-lesioned participants, suggesting these behaviors depend on PRC and not hippocampal structures. We note that these results are consistent across all computational models evaluated (Supplemental Fig. S3).

There are numerous limitations in the retrospective analysis, some inherited from the literature itself. First, there are statistical concerns. Low sample size in the retrospective dataset is a consequence of the number of (total and available) experiments in the literature. Moreoever, the available experiments did not estimate the participant- or item-level reliability. Second, there is considerable off-hypothesis variance in experimental designs across studies. While the minimal number of objects required for an oddity trial is three, experiments vary in number of objects presented, ranging from three to seven (Supplemental Fig. S2 and S4). Our modeling approach is agnostic to the task demands introduced by changes in the number of objects to be discriminated between, but we cannot say the same for human participants, where objects may increase extra-perceptual (e.g. memory-related) task demands. Third, while our results suggest PRC-lesioned performance relies on high-level visual cortex, the available stimuli do not enable us to clearly isolate early-from late-stage processing within the VVS (Fig. 3). Finally, our retrospective analysis relates human behavior to a computational proxy for the VVS, not directly to human VVS neural responses.

We address many of these limitations within the retrospective experiment with a novel oddity experiment (Fig. 4) where we directly compare PRC-intact human performance to electrophysiological recordings collected from the macaque VVS (Majaj et al., 2015). Unlike results from the retrospective dataset, the novel dataset clearly separates V4-from IT-supported performance (Fig. 5a). As predicted, model performance approximates IT-supported performance (Fig. 3b). Critically, PRC-intact behavior substantially outperforms IT-supported performance (Fig. 5c). Moreover, we find that the divergence between PRC-intact and IT-supported behavior is reliably predicted by human reaction time: On an item-by-item basis, the degree to which PRC-intact performance exceeds IT-supported performance scales linearly with the time required (Fig. 5d). A computational proxy for high-level visual cortex (but not a linear readout of V4) exhibits the full pattern of accuracy and reaction time data relating PRC-intact to IT-supported performance, further validating our use of the model in the retrospective analysis. Data from this novel experiment confirm predictions central to the perceptual-mnemonic hypothesis with unprecedented resolution and validate previous computational results. Additionally, these data offer the first account of the temporal demands inherent in ostensibly PRC-dependent visual discrimination behaviors: supra-VVS performance scales linearly with reaction time.

Results from our retrospective and novel analyses validate core tenets of the perceptual-mnemonic hypothesis: PRC provides an advantage in visual discrimination tasks beyond VVS-supported performance, PRC’s contribution to these behaviors is independent of the hippocampus, and PRC-lesioned performance appears to rely on the VVS. We make no claim as to whether these PRC-dependent behaviors should be described as either ‘perceptual’ or ‘mnemonic.’ In fact, it may be that these psychological constructs are not well suited to characterize PRC-dependent computations, or neural function more generally (Jolly and Chang, 2019). Indeed, the retrospective analysis offers a case study on the limitations of such an approach: Experimentalists *were* able to design incisive tests of PRC function by operationalizing constructs such as stimulus ‘complexity,’ but an equal number of non-diagnostic stimulus sets emerged, on both sides of the debate, without any measurable way to disambiguate between competing interpretations of the available data. Our approach offers an extensible alternative, enabling us to situate lesion, electrophysiological, and behavioral data within a unified theoretical framework. This work underscores the promise of developing computational models of PRC that operate over experimental stimuli, as they are presented to experimental participants. We believe that this stimulus-computable, biologically plausible modeling approach will continue to offer novel insights into how the MTL supports such enchanting—indeed, at times, indescribable—behaviors.

## 4 Acknowledgements

This work is supported by a National Science Foundation Graduate Research Fellowship under Grant No. DGE–1656518, Stanford’s Center for Mind Brain Behavior and Technology, and the Marcus and Amelia Wallenberg Foundation (MAW2015.0043). We thank the original authors of the studies in the retrospective dataset, who generously provided stimuli when possible, and their assistance even when the stimuli were not accessible. Specifically, we would like to thank Morgan Barense, Elizabeth Buffalo, Tim Bussey, Lila Davachi, Andy Lee, Elizabeth Murray, Larry Squire, and Craig Stark, as well as Jennifer Frascino and Mona Hopkins for their diligent efforts securing multiple stimulus sets. We thank Akshay Jagadesh for insightful conversations and suggestions throughout the course of this work, as well as Mark Eldridge, Chiara Giovanni, Nathan Kong, Heather Kosakowski, Emily Mackevicius, Ayesha Nadiadwala, Russell Poldrack, and Natalia Vel’
sez for their feedback on previous versions of this manuscript. Finally, we thank the human and non-human participants in this research, without whom this work would not have been possible.

## 5 Author contributions

T.B. and A.D.W. conducted the literature review and reached out to original authors. T.B. conceived of the modeling approach and performed all modeling work, under the supervision of D.L.K.Y.. T.B. designed, implemented, and analyzed the novel experiments. T.B., D.L.K.Y., and A.W.D. discussed results, then wrote and revised the manuscript.

## 6 Declaration of Interests

The authors declare no competing interests.

## 7 Inclusion and Diversity

One or more of the authors of this paper belongs to a racial group that is underrepresented in the sciences. One or more of the authors of this paper received funding from a program designed to support underrepresented communities in the sciences. While citing references scientifically relevant for this work, we also actively worked to promote gender balance in our reference list.

## STAR METHODS

### RESOURCE AVAILABILITY

#### Lead Contact

Further information and requests for stimuli, models, and analysis procedures should be directed to and will be fulfilled by the lead contact, Tyler Bonnen.

#### Materials Available

This study did not generate new unique reagents.

#### Data and Code Availability

The online repository at https://github.com/neuroailab/mtlperception.git contains all models, statistical analyses, and visualizations used within the current study.

### EXPERIMENTAL MODEL AND SUBJECT DETAILS

A total of 297 human participants completed the novel visual discrimination task (Results: Novel Dataset) via on online crowd-sourcing platform (Amazon Mechanical Turk). All participants provided informed consent in accordance with procedures approved by Stanford’s Institutional Review Board.

### METHODS DETAILS

#### Literature Review

Criteria for inclusion in the retrospective analysis were threefold. First, the data must include behavior from both PRC-lesioned and -intact participants. Second, the experiment must have been administered to either human or non-human primate participants. Third, participants must have performed oddity visual discrimination tasks. The initial Google Scholar search terms used were “perirhinal lesion oddity” resulting in 425 results. The terms ‘human’ or ‘primate’ were not included in this search as experimental participants in human research are often referred to simply as ‘subjects.’ After candidate experiments were identified from these 425 results, the references cited in each of these candidate papers were used as a source of candidate papers missed in the initial search. An additional exclusion criterion was incorporated, as one oddity experiment (Lee & Rudebeck 2010) required that participants reference real-world shape properties of objects not presented on the stimulus screen alongside the stimuli. This experiment was not included in further analysis. The corresponding authors in each experiment were contacted via email and asked to provide experimental materials necessary for the current computational approach. This included: (a) First, behavioral data from PRC-lesioned and -intact participants with the finest granularity that could be collected (e.g. trial, subject, or group level data). When available, this also included behavioral data from hippocampal-lesioned and hippocampal-intact participants. (b) Second, the stimuli corresponding to these behavioral data—ideally, the exact stimuli presented in each experiment conducted. For all studies, the corresponding authors (or their associates) responded promptly and were eager to provide the data requested. The complete list of studies identified through this search is as follows: Barense et al., 2007; Buckley et al., 2001; Buffalo et al., 1998a; Inhoff et al., 2019; Knutson et al., 2012; Lee et al., 2006; Lee et al., 2005a; Lee et al., 2005b; Levy et al., 2005; Shrager et al., 2006; Stark and Squire, 2000.

#### Retrospective Dataset

The stimuli and behavioral data were were able to obtain only included human participants. Across all of the obtained experiments, we were able to reliably secure experiment-level behavioral data (i.e. averaged across trials) for each group within a given study (e.g. the performance of PRC-lesioned participants performing condition A, B, etc., within a given study). In order to compare model and human behaviors, we compare behavior at the level of the experiment (i.e. averaged across trials). For most of the obtained experiments, the exact trial-level stimuli presented to participants were used in the modeling approach. However, there were two experiments (Stark and Squire, 2000 and Lee et al., 2006) where the distribution of stimuli presented in each experiment had to be approximated. For Stark and Squire, 2000, the authors randomly selected stimuli to be used in each trial, from a set of all possible stimuli. They could not recover the exact trial-by-trial stimuli shown to experimental participants. Instead, the corresponding authors provided all stimuli used across faces and “snow” (partially occluded object) experiments, as well as the pseudo-random protocol used to generate each experiment: For each “typical” item, five different viewpoints were drawn from all available stimuli of this item. Faces had a total of six items, each corresponded to different (but common across faces) viewpoints. For each object, there were a total of five viewpoints, such that all viewpoints of this item were used in each trial. In ‘snow’ conditions, for each trial, the typical object was selected at random, and all of its exemplars were used; the oddity object was selected at random, and one of its exemplars was selected at random to be that trial’s oddity. For faces, after selecting a typical face, and a subset of 5 of its exemplars, the oddity identity was sampled randomly, with a viewpoint distinct from that present in the typical faces. Consequently, each face trial included an oddity that was always from a different viewpoint from all typical faces. For Lee et al., 2006, the corresponding authors were able to provide all stimuli. However, as with Stark and Squire, 2000, in experiment two only a subset of the stimuli were presented to participants. Across participants, the number of trials in this subset was constant (31/40), but the exact items presented to each subject was drawn randomly from all available stimuli. For both the Stark and Squire, 2000 and Lee et al., 2006 we approximate the stimuli presented to participants by generating a population of experiments (N=100) that adhered to the protocols outlined above. We then compare the model performance across this population of experiments (i.e. averaged performance across all N iterations generated by this sampling protocol) to the obtained human behavior for each experiment.

#### Model Fit to Electrophysiological Data

We use one instance of a task-optimized convolutional neural network (VGG16, Simonyan and Zisserman, 2015), implemented in tensorflow and pre-trained to perform object classification on a large-scale object classification dataset (Deng et al., 2009). To identify a model layer that best fits IT cortex, we utilize previously collected (Majaj et al., 2015) electrophysiological responses from macaque V4 and IT cortex, along with the stimuli that elicited these responses. Using ‘medium’ and ‘high’ variation images from this data set, we convert each image from greyscale to RGB then resize it to accommodate model input dimensions (224×224×3). We pass each image to the model and extract responses from all layers (e.g. convolutional, pooling, and fully connected layers), and vectorize each layer’s output. We randomly segment these model responses to each image into training and testing data using a 3/4th split. Thus, we have multi-electrode responses from macaque V4 and IT to a set of images, alongside model responses to those same images. For each layer, we learn a linear mapping between vectorized model responses and a single electrode’s responses to the training images, using sklearn’s implementation of PLS regression (with five components). We evaluate this mapping between model and neural responses by computing the Pearson’s correlation between model-predicted responses and observed responses for each electrode across all test images. For each layer, this results in a single correlation value for each electrode, which we repeat over all electrodes. This results in a distribution corresponding to that layer’s cross-validated fits to population-level neural responses, both for electrodes in IT and V4. We compute the split half reliability for V4 (*r* = .63 ± .22STD) and IT (*r* = .73 ± .24STD) across neurons in each region. We then divide the distribution of cross-validated fits to IT and V4 by the reliability in each region—as a noise-corrected adjustment. This results in a single score—the noise-corrected, median cross-validated fit to both IT and V4—which we repeat across all layers (Fig. 3a: black and dotted lines for IT and V4 fits across layers, respectively). We determine also each layer’s differential fit with primate IT (Δ_*IT-V*4_) by taking the difference between the model’s fit to IT and V4 (Fig. 3a: hollow line). Early model layers (i.e. first half of model layers) better predict neural responses in early (V4) regions of the visual system (ttest *t*(8) = 2.70, *P* = .015), with peak V4 fits occuring in pool3 (noise-corrected *r* = .95 ± .30STD) while later layers (e.g. second half of model layers) better predict neural responses in more anterior (IT) regions (ttest *t*(8) = 3.70, *P* = .002), with peak IT fits occuring in con5 1 (noise-corrected *r* = .88 ± .16STD). We use model responses at this layer, con5 1, as an ‘IT-like’ model layer in subsequent analyses.

#### Model Performance on Retrospective Dataset

For each trial, in each available experiment, the stimulus screen containing N objects was segmented into N object-centered images, using one of three protocols. For some experiments (e.g. Stark and Squire, 2000), stimuli were already segmented, requiring no additional processing. For other experiments (e.g. Lee et al., 2006), the stimuli for each trial were only available in the form presented to experimental participants, as a single image containing multiple objects. For these experiments, for each trial, the stimulus screen was segmented using a kmeans clustering approach that automatically identified the centroid of each object, defined a bounding box around each of these centroids, and extracted each object from the coordinates of each bounding box. There was a final class of experiments with more irregular dimensions (e.g. ‘Familiar’ objects in Barense et al., 2007; these stimuli were segmented by splitting the original stimulus screen into quadrants of equal size. Additionally, these images were rotated (by 90, 180, or 270 degrees) such that each object was presented as close to its canonical viewpoint as possible. For all segmentation protocols, we passed each trial’s N object-centered images to the model, then extracted model responses from an ‘IT like’ layer. These layer responses were flattened into length F vectors, resulting in an FxN response matrix for each trail. To identify the item-by-item similarity between objects in this trial, were used Pearson’s correlation between items in this FxN response matrix, generating an NxN (item-by-item) correlation matrix. The item with the lowest mean off-diagonal correlation was the model-selected oddity (i.e. the item least like the others) which we then labeled as either correct or incorrect, depending on its correspondence with ground truth. After repeating this protocol (for visualization see Fig. 1c) for each trial in the experiment, we computed the average accuracy across all trials. This single value, ‘model performance’, represents the performance that would be expected from a uniform readout of IT.

#### Non-Diagnostic Experiments

By definition, experiments that are fully supported by canonical VVS regions are not diagnostic of PRC involvement in perception; if the VVS enables 100% accuracy on a given experiment, no further perceptual processing should be necessary. This does not, however, imply that human performance on these VVS-supported tasks will also be at ceiling: While a *lossless* readout of the VVS should perform these tasks at ceiling, a *lossy* readout—due to, for example, attentional demands of maintaining those perceptual representations between saccades—may be below ceiling. In this way, below-VVS performance on these trials can be attributed to extra-perceptual task demands that are orthogonal to the perceptual-mnemonic hypothesis. As a validation, we observe that all color experiments in the retrospective dataset adhere to this logic: Model performance achieves 100% accuracy on all trials (both ‘Easy’ and ‘Difficult’ experiments) and PRC-lesioned performance on these conditions is statistically indistinguishable from PRC-intact behavior (Barense et al., 2007). Nonetheless, human performance on ‘difficult’ trials is significantly lower than ‘easy’ trials. These results corroborate researchers’ expectations that these control stimuli are not diagnostic of PRC function, while the difficulty manipulation imposes extra-perceptual task demands. We use this logic to identify experiments that are not diagnostic of PRC involvement in perception.

First, we estimate model performance for all experiments in the retrospective dataset. We then identify those stimulus sets where model performance is 100% accurate. As expected, this identifies control experiments (e.g. color experiment in Barense et al., 2007). Interestingly, this also identifies many experiments that the original authors described as ‘complex,’ and subsequently used to evaluate the role of PRC in perception. These experiments belong to two groups. The first group contains experiments that were argued as evidence *against* perirhinal involvement in perception (Buffalo et al., 1998a; Knutson et al., 2012) because performance did not significantly differ between PRC-intact and -lesioned participants. However, model performance suggests that canonical VVS regions should be sufficient for ceiling performance (Supplemental Fig. S1a-b; top row); consequently, the matched PRC-lesion/intact performance is expected, and entirely consistent with predictions from the perceptual-mnemonic hypothesis. The second group contains experiments that were argued to reveal evidence *in support* of perirhinal involvement in perception (Barense et al., 2007; Inhoff et al., 2019) because PRC-lesioned subject behavior was impaired relative to PRC-intact controls. However, the model suggests that canonical VVS regions should be entirely sufficient for performance on these tasks (Supplemental Fig. S1c-d; top row); consequently, the observed divergence may not be due to perceptual demands in these tasks.

We note that these same experiments are identified as non-diagnostic using multiple model-based and model-agnostic protocols. First, using pytorch’s implementations (Paszke et al., 2017) of multiple instances within this convolutional model class (e.g. Densenet 201, Huang et al., 2017, Resnets, He et al., 2016, Alexnet, Krizhevsky et al., 2012, Squeezenet, Iandola et al., 2016), we replicate the analyses above: For each model, we estimate model performance from an IT-like layer. All models evaluated identify the same 14 experiments as non-diagnostic. Next, we repeat this approach using randomly initialized (i.e. untrained) models. Again, we find that the same experiments are identified as non-diagnostic, suggesting that object-specific training is not necessary for ceiling performance. Next, we repeat the analysis above over pixel-level representations, forgoing the use of the model entirely. For each object in a trial containing *N* objects, we flatten the original images, converting them into *N* vectors. We then compute the pairwise correlation between these vectors, generating an *N* by *N* covariance matrix. As in the model-based protocol, we identify the item with the lowest off-diagonal correlation as the oddity. That is, the decision making protocol is unchanged, but it operates on the pixel-level covariance matrix. This approach again achieves 100% accuracy on the same 14 experiments, and only these 14 experiments: these experiments can be solved by determining *pixel-level* differences between objects. Not only do all computational proxies of the VVS achieve 100% accuracy on these experiments, even untrained models and pixel-level analyses consistently identify the same set of 14 experiments as non-diagnostic.

After excluding these non-diagnostic experiments, there are 14 experiments that are positioned to evaluate the involvement of PRC in perception. This includes 10 experiments the original authors identified as diagnostic (all ‘snow’ experiments in Stark and Squire, 2000, ‘high ambiguity’ experiments, both ‘novel’ and ‘familiar’ experiments in Barense et al., 2007, ‘novel objects’ and ‘faces’ experiments in Lee et al., 2005a, and ‘different faces’ experiments in Lee et al 2006). Additionally, this includes 4 experiments that were designated as ‘control trials’ by the original authors (‘low ambiguity novel objects’ and ‘low ambiguity familiar objects’ in Barense et al., 2007, ‘familiar objects’ in Lee et al., 2005a, and ‘different scenes’ in Lee et al., 2006). Note that the only criteria for this analysis is that model performance is not at ceiling: This selection procedure makes no claim about whether each individual experiment will exhibit PRC-related deficits. We refer to these as ‘diagnostic’ experiments throughout the Results and Discussion.

#### VVS Reliance

As outlined (STAR Methods: Model Fit to Electrophysiological Data), we estimate each model layer’s fit to electrophysiological responses in macaque IT and V4 (Fig. 3a: solid black, dashed lines for IT, V4, respectively). Here we compute each layer’s differential fit to IT by taking the difference between noise-corrected IT and V4 fits (Fig. 3a: Δ_*IT-V*4_, hollow). Differential fit to IT cortex increases in ‘deeper’ layers (*β* = .98, *F* (1, 17) = 21.75, *P* = 10^*-*13^).

Next, we determine each layer’s fit to human behavior in the retrospective dataset using the mean squared prediction error (MSPE) between human and model behavior, for each subject group: 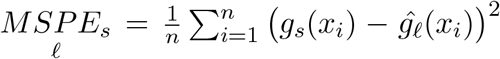 where *x*_*i*_ is each experiment, ℓ is a single layer within the 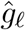 is the protocol used to estimate model performance that operates over all trials in *x*_*i*_. This results in estimates for model performance on each experiment, for each layer of the model. We compare this layer-by-experiment estimate of model performance with the performance of participants in group *s* on experiment *x*_*i*_, *g*_*s*_(*x*_*i*_). We determine the difference between model 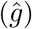 and human (*g*) performance across all experiments, and then average, resulting in a single value for the fit to each subject group *s*, from each model layer (e.g. *MSPE*_*prc*.*lesion*_). Finally, we compute the relative fit between lesioned and intact subjects by computing the difference between model fits to lesioned and intact humans, Δ_*group*_ = *MSPE*_*intact*_ - *MSPE*_*lesion*_ for both PRC- and HPC-lesioned groups (e.g. Δ_*prc*_ = *MSPE*_*prc*.*intact*_ - *MSPE*_*prc*.*lesion*_).

To assess whether PRC-lesioned behavior is better fit by late-stage processing within the VVS we relate the model’s relative fit with lesioned performance (for both Δ_*prc*_ and Δ_*hpc*_) to the model’s differential fit to IT cortex (Δ_*IT-V*4_). Model layers that better fit IT cortex (Δ_*IT-V*4_) are better predictors of relative fit to PRC-lesioned behavior (Δ_*prc*_, Fig. 3c: top). Additionally, we determine whether the interaction between lesioned and intact subject behavior is significant, repeating previous analyses across all layers, for each patient group. Only ‘IT-like’ layers demonstrate significant interactions between subject groups (e.g. PRC-lesioned vs PRC-intact) after correcting for multiple comparisons across layers. There is no correspondence with HPC-lesioned behavior (Δ_*hpc*_, Fig. 3c: bottom).

#### Novel Stimulus Set Generation

We utilize stimuli and electrophysiological data from a previous experiment (Majaj et al., 2015) consisting of 5760 unique images, each with population-level electrophysiological responses recorded from primate V4 and IT. Every black and white image contains one of 64 objects, each belonging to one of eight categories, rendered in different orientations and projected onto random backgrounds— for a total of 90 images per object. We reconfigure these stimuli into within-category oddity tasks. Each trial is designed to have the minimal configuration of objects (*n* = 3) required to be an oddity task: two of the three objects share an identity (two images of the ‘typical’ object_*i*_, presented from two different viewpoints and projected onto different random backgrounds) and the other is of a different identity (one image of the ‘oddity’, object_*j*_, e.g. two animals, where ‘elephant’ and ‘hedghog’ are object_*i*_ and object_*j*_, respectively). We generate a sample trial_*ij*_ for the pair_*ij*_ of objects *i* and *j* by randomly sampling two different objects from the same category, then sampling two images of object_*i*_ (without replacement) and one image of the oddity of object_*j*_, all with random orientations and backgrounds. These three images comprise sample_*ij*_ of the pair_*ij*_.

#### Model Performance on Novel Stimuli & Comparing Readout Performance

We first determine model performance on our novel stimuli using a uniform, linear readout of model responses (i.e. the distance metric used in the retrospective analyses). For each object pair_*ij*_, we generate 100 random sample_*ij*_s, pass each object through the model, extract model responses from an IT-like layer, and determine the item with the lowest off-diagonal correlation as the model-selected oddity. We average across these binarized (100 correct or incorrect) outcomes for each randomly generated sample_*ij*_, which we repeat over all pairs. This serves as a uniform, linear measure of model performance (model performance_*uniform*_). We order the results according to performance, compute the difference between each adjacent pair_*ij*_ (Δ_*pair*_) and note that these 448 unique pairs (Fig. S5a: black) densely and continuously span the range of model performance (averaged 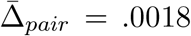). We select four categories that continuously span the space of model performance (*min* = .26, *max* = 1.0, 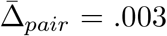 Fig. S5b: Faces, Chairs, Planes, and Animals), which contains a total of 224 unique typical-oddity pairs and use these objects for the human psychophysics experiments.

Next, we develop a more expressive estimate of model performance using a weighted, linear readout of model responses. We use a modified leave-one-out cross validation strategy to develop this approach. For a given sample_*ij*_ trial we construct a random combination of three-way oddity tasks to be used as training data; we sample without replacement from the pool of all images of object_*i*_ and object_*j*_, excluding only those three stimuli that were present in sample_*ij*_. This yields ‘pseudo oddity experiments’ where each trial contains two typical objects and one oddity that have the same identity as the objects in sample_*ij*_ and are randomly configured (different viewpoints, different backgrounds, different orders). These ‘pseudo oddity experiments’ are used as training data. We reshape all images, present them to the model independently, and extract model responses from an ‘IT-like’ model layer (in this case, we use fc6 which has a similar fit to IT as conv5 1 but fewer parameters to fit in subsequent steps). From these model responses, we train an L2 regularized linear classifier to identify the oddity across all (*N* = 52) trials in this permutation of pseudo oddity experiments generated for sample_*ij*_. After learning this weighted, linear readout, we evaluate the classifier on the model responses to sample_*ij*_. This results in a prediction which is binarized into a single outcome {0 | 1}, either correct or incorrect. We repeat this protocol across 100 random sample_*ij*_s, and average across them, resulting in a single estimate of model performance for each pair_*ij*_ (model performance_*weighted*_).

Finally, we compare model performance from these uniform and weighted approaches—two linear readouts of model performance. As expected, the weighted readout of model responses outperforms a uniform distance metric (paired ttest, *t*(447) = 33.55, *P* = 10^*-*123^; Supplemental Fig. S5a: points on the y axis consistently above the diagonal). Nonetheless, there is a linear relationship between uniform and weighted estimates of model performance (ols regression *β* = 1.01, *F* (1, 446) = 23.28, *P* = 10^*-*79^). We refer to this expected value for model performance_*weighted*_ given our observed model performance_*uniform*_ as model performance_*transformed*_. We can use this transformation to estimate the performance that would have been observed if we were able to use a weighted readout in the retrospective analysis, projecting model performance_*uniform*_ into model performance_*transformed*_, which approximates model performance_*weighted*_. This transformed model performance_*transformed*_ does significantly better at predicting PRC-lesioned behavior than the original model performance_*uniform*_ (interaction between readout types in predicting PRC-lesioned performance *β* = -.20, *F* (2, 25) = -4.26, *P* = 2×10^*-*4^; Supplemental Fig. S5b), to the degree that there is no longer a significant difference between PRC-lesioned performance and model performance. This improved fit to PRC-lesioned performance motivates the need for future work that estimates model performance using weighted readouts. We use our weighted estimate of model performance_*weighted*_ as the default estimate of model performance in the novel experiment.

#### High-throughput Psychophysics Experiments

We create oddity trials composed of stimuli containing these 224 objects identified in the preceding analyses. To create each trial, we adopt the same the protocol used to generate each sample_*ij*_. We use this protocol for each of the 224 pair_*ij*_s: we generate 5 random combination of trials from each pair_*ij*_ and fix these trials across all experiments (i.e. trial_*ij*1_, trial_*ij*2_, …, trial_*ij*5_), resulting in (224 × 5) 1120 unique trials. We administer a randomized subset (*N* = 100) of these oddity trials to 297 human participants using a browser-based experimental paradigm implemented in jsPsych (De Leeuw, 2015).

Each trial is self paced, such that participants initiate the beginning of the trial with a button press (spacebar). In each trial, one of 1120 oddity stimuli is presented for 10 seconds. Participants are free to respond at any point with a button press, indicating the location of the oddity they have identified (right, left, bottom). If participants respond before the trial ends, their responses are recorded, and the trial is terminated. If a participant fails to respond within 10 seconds, the trial is terminated and marked as incorrect. After an initial familiarization phase (5 trials) to acclimate participants to the task, no further feedback is given at any point during the experiment. All participants are compensated with a initial base rate. Additionally, each subject is given a monetary bonus for each correct answer, or a monetary penalty for each incorrect answer. This monetary incentive structure was titrated to ensure that participants are encouraged to attempt even the most difficult perceptual trials, while ensuring that all participants are compensated fairly (at least earning California’s minimum wage for the time they participate in the experiment). At the end of each experiment, participants are informed of their performance, alongside their total bonus. All tasks are completed through Amazon’s Mechanical Turk.

Given the truly random experimental generation protocol—and, subsequently, the highly variable nature of the stimuli that comprise each trial—there is no guarantee that a given trial_*ijn*_ will contain the information sufficient for a correct response; for example, all the faces in a trial might be rotated out of view, such that the correct oddity can not be determined. To address this, of the 5 stimuli presented, for each of the 224 pair_*ij*_s, we restrict our analysis to 1 trial_*ij*_ which is identified empirically. We select this exemplar for each pair_*ij*_ using a single criterion: the item whose average accuracy (across participants) is closest to the average accuracy measured across all trials (across participants) belonging to other categories. This procedure enables us to exclude outliers (due to, for example, the objects not being fully visible on the viewing screen) while not biasing the results in future analyses. We note, however, that all results from this dataset are qualitatively identical if we run these analyses over all trials, not simply over the exemplar for each pair_*ij*_. Regardless, for all analyses, performance estimates are computed across the population of human participants. In this pooled population of behavior, we compute the reliability by determining the averaged correlation over 1000 randomly shuffled split halves: accuracy was reliable at the category (averaging across all oddity trials in a given category, *r* = .97 ± .03), object (averaging over all oddity objects for a given typical object, *r* = .69 ± .07), and image level (average performance on a given trial, averaged across participants, *r* = .24 ± .05). This effect was even more prominent in the estimates of reaction time at the category (*r* = .99 ± .01), object (*r* = .91 ± .02), and image level (*r* = .76 ± .02).

To relate human performance on these oddity tasks with model performance_*weighted*_, we employ the same pseudo experimental leave-one-out cross-validation strategy as outlined above, but with 100 train-test splits for each trial_*ij*_, across all (*N* = 224) unique typical-oddity pairings. To relate human (and model) performance with the electrophysiological data, we repeat the leave-one-out cross-validation strategy developed for determining model performance, but in place of the fc6 model representations, we run the same protocol on the population level neural responses from IT and V4 cortex. We perform all analyses comparing human, electrophysiological, and model performance at the object level: for each object_*i*_ we average the performance on this object across all oddities (i.e. object_*j*_, object_*k*_, …) resulting in a single estimate of performance on this item across all oddity tasks, for each (*N* = 32) item.

### QUANTIFICATION AND STATISTICAL ANALYSIS

#### Model Fit to Electrophysiological Data

We split all ‘high’ and ‘medium’ variation images (*n* = 3840) into a training (*n* = 2880) and testing (*n* = 960) set, using a random 3/4th split. Given the training images, we learn a linear mapping between model responses and electrode responses, independently for each electrode (*n*_*electrodes*_ = 296), using sklearn’s implementation of PLS regression (with five components). We evaluate this model-electrode mapping by computing the Pearson’s correlation between model-predicted responses and observed responses across all test images, for each electrode, for each layer. We partition these electrode responses by region, evaluating model-by-layer fit to electrodes in neural area V4 (*n* = 128) and inferior temporal (IT) cortex (*n* = 168). This yields a distribution corresponding to that layer’s cross-validated fits to population-level neural responses, both from electrodes in IT and V4. We compute the split-half reliability for V4 and IT across neurons in each region, using 1000 train-test splits. We then use this split-half reliability for neurons in each region as the noise ceiling. For each layer, we divide the distribution of cross-validated fits to IT and V4 by the reliability in each region as a noise-correcting adjustment. This results in a single score for each layer’s fit to each brain region: the noise-corrected, median cross-validated fit to both IT and V4 (Fig. 3a: black and dotted lines for IT and V4 fits across layers, respectively). We determine also each layer’s differential fit with primate IT, Δ_*IT-V*4_, by taking the difference between the model’s fit to IT and V4 (Fig. 3a: hollow line). We identify the most ‘IT-like’ model layer using results reported in Fig. 3a, Results, and STAR Methods: Methods Details.

#### Retrospective Dataset

To estimate ‘model performance’ across all experiments within the retrospective dataset we extract trial-level model responses from an ‘IT-like’ layer. With these model responses, we identify the model-selected oddity as the item which has the lowest between-object correlation in that trial, using python’s implementation of correlation coefficients within numpy (https://numpy.org/). To estimate model performance we simply average across all trials in a given experiment. Within the retrospective dataset, we identify experiments (*n* = 14) that can be used to evaluate the involvement of PRC in perception. We use python’s statsmodels (https://www.statsmodels.org/) implementation of parametric statistical tests to evaluate the relationship between PRC-intact, -lesioned, and model performance across these experiments. We first report the outcomes of a paired t-test comparing the relationship between PRC-intact and -lesioned performance (Results: PRC-lesioned subjects are impaired on oddity tasks). We then compare PRC-intact, -lesioned, and model performance across these experiments with an interaction model (i.e. ‘human performance ∼ model performance * lesion group’) using statsmodels implementation of OLS regression (Results: A computational model of the VVS approximates PRC-lesioned performance, and Fig. 2). Finally, we use the same statsmodels implementation of OLS regression to relate PRC-intact and-lesioned performance to model performance estimated from all layers (Results: PRC-lesioned oddity performance appears to rely on high-level visual cortex, STAR Methods: VVS Reliance, and Fig. 3).

#### Novel Stimulus Set

We determine model performance from an ‘IT-like’ layer on each trial within our novel stimuli using two methods. First, the distance-based metric from the retrospective analyses. Second, by using a weighted (i.e. learned), linear readout of model responses, using sklearn’s implementation of L2 regularized logistic regression (https://scikit-learn.org/). We use a modified leave-one-out cross validation strategy to develop this approach, repeating this protocol across 100 random permutations for each trial to get a stable estimate of the trial-level performance. We compare the two methods using the statsmodels implementation of OLS regression. We repeat this approach using electrophysiological data to estimate V4- and IT-supported performance. Finally, we estimate human performance on these same stimuli: estimates are computed across the population of human participants (*n* = 297). We can compare these estimates (V4-supported, IT-supported, and model performance) to the performance of human participants’ discrimination between these same stimuli. We perform all analyses comparing human, electrophysiological, and model performance at the object level: for each object_*i*_ we average the performance on this object across all oddities (i.e. object_*j*_, object_*k*_, etc.) resulting in a single estimate of performance on each item (*N* = 32) across all oddities. We use python’s statsmodels implementation of OLS regression to evaluate the relationship between V4-supported, IT-supported, model performance, and human performance (Results: PRC-intact participants outperform electrophysiological recordings from IT, Results: Model Performance on Novel Stimuli & Comparing Readout Performance, Fig. 5).

## Supplemental Information

**Supplementary Figure S1:**
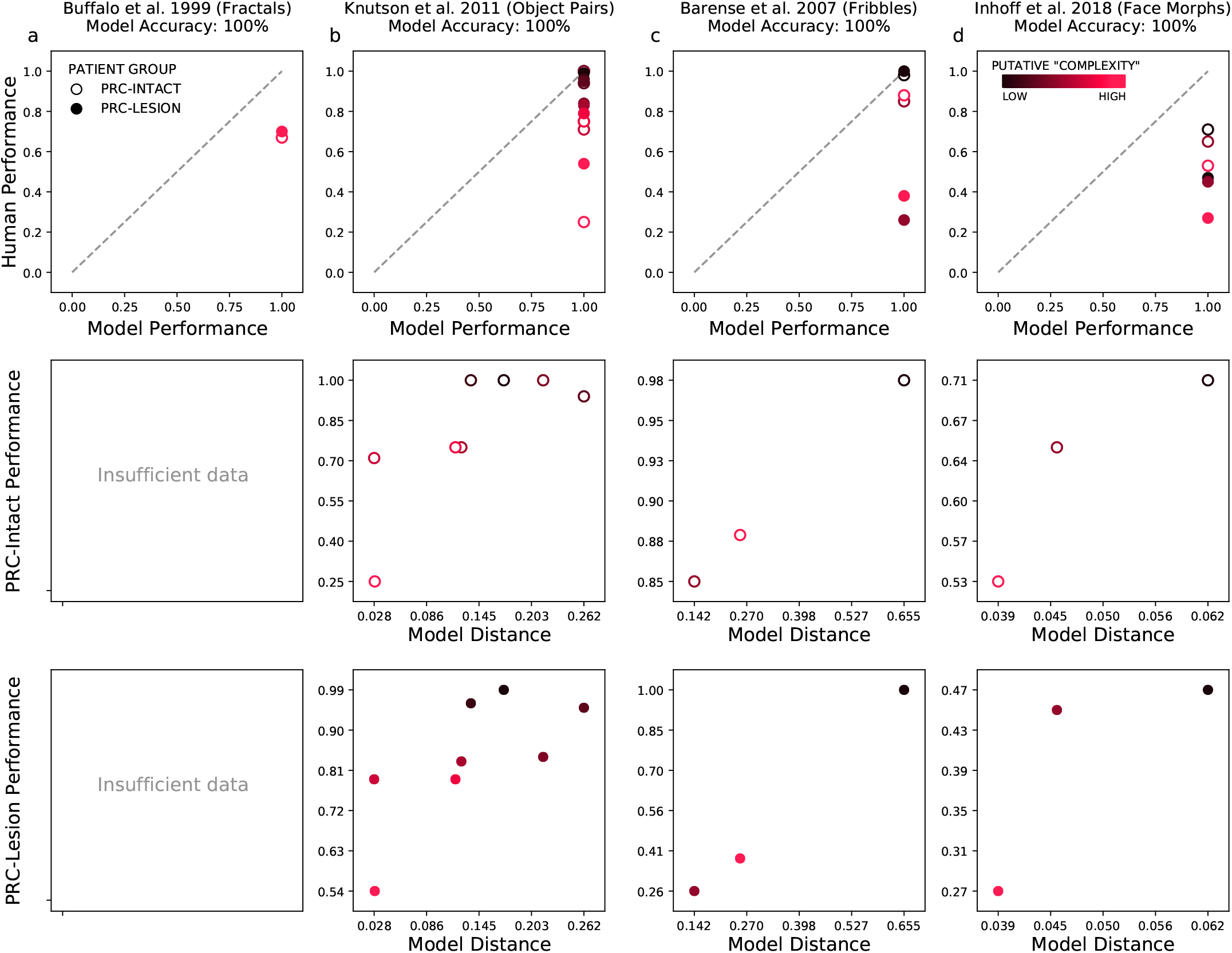
Non-diagnostic experiments exist on both sides of the debate, Related to Figure 2. We find that numerous experiments used to evaluate PRC involvement in perception are perfectly supported by a computational proxy for the VVS: Model performance is 100% from an IT-like layer (**row one**). Results from these non-diagnostic experiments belong to two categories: eight experiments used as evidence against perirhinal involvement in perception, as performance did not significantly differ between PRC-intact and -lesioned participants (a-b; all experiments in Knutson et al., 2012 and Buffalo et al., 1998a), and six experiments used as evidence in support of perirhinal involvement in perception, as PRC-intact performance was significantly impaired relative to their PRC-intact counterparts (c-d; all ‘Fribbles’ in Barense et al., 2007, all stimuli in Inhoff et al., 2019). While our approach is not designed to account for behavior collected from these non-diagnostic experiments, we can nonetheless provide a *descriptive* account of our modeling results that is consistent with the perceptual-mnemonic hypothesis. For non-diagnostic tasks that have been used as evidence against PRC-involvement in perception (a-b), PRC-lesioned subjects demonstrate no impairments because these tasks can be fully supported by the VVS. As such, no PRC-related perceptual deficits are expected. Meanwhile, on non-diagnostic tasks used as evidence in support of PRC-involvement in perception (c-d), PRC-lesioned impairments may instead be due to extra-perceptual task demands (e.g. mnemonic demands imposed by the relatively large number of objects per trial in ‘Fribbles’ experiments) or other confounding factors, such as the extent of lesion in PRC-adjacent cortical structures. This is not evidence against the perceptual-mnemonic hypothesis, but simply that these stimuli are not well positioned to evaluate its central claims. Next, we relate PRC-lesioned and -intact performance in experiments a-d to a different model readout: model distance (**row two and three**). Here, we estimate the averaged distance (1 - *r*) between the oddity and typical objects across trials in each experiment, which we refer to as ‘model distance.’ While low experimental *N* s warrant caution interpreting these data, within each study model distance predicts performance of both PRC-intact (row two) and -lesioned (row three) participants. Specifically, participants demonstrate higher accuracy for experiments associated with greater model distance. While model performance enables us to exclude non-diagnostic experiments from the retrospective dataset, using a different model readout for the experiments nonetheless predicts the relative performance of human participants in these experiments. We note that the × and y axes are rescaled across columns; the magnitude of these effects are not common across studies. Nonetheless, these results are consistent with expectations that below-ceiling performance on VVS-supported experiments is due to extra-perceptual task demands, where increasing similarity between items results in greater demands on extra-perceptual cognitive systems.

**Figure S2:**
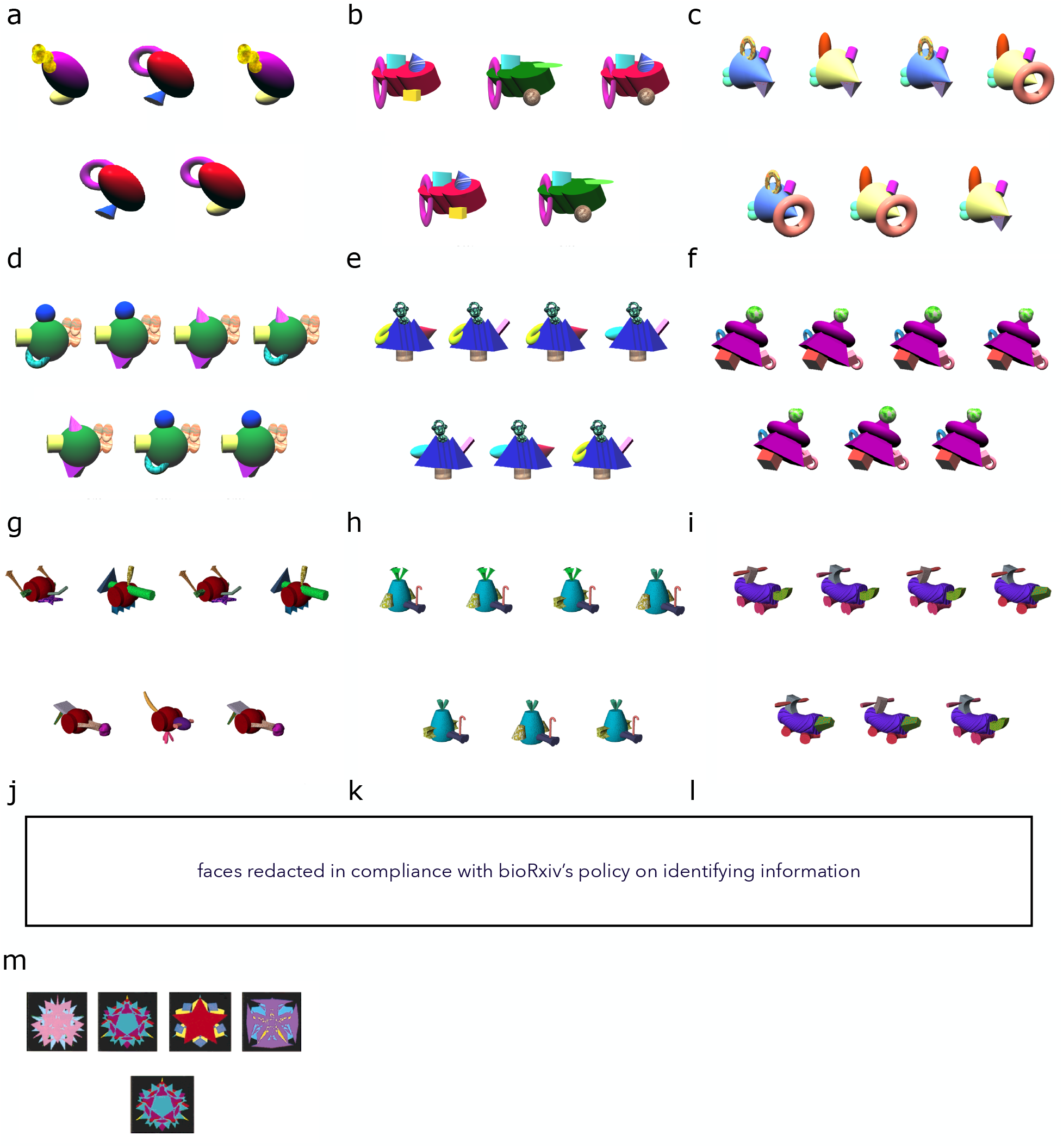
An example trial from each experiment identified as non-diagnostic, Related to Figure 2. All non-diagnostic experiments (i.e. all experiments on which model performance achieves 100% accuracy) contain objects presented from the same viewpoint. We include example trials for each experiment within each study: **(a-f)** Knutson et al., 2012 difficulty conditions 1-6; **(g-i)** low, medium, and high ambiguity ‘Fribbles’ in Barense et al., 2007; **(j-l)** easy, medium, and hard conditions in Inhoff et al., 2019; **(m)** complex stimuli in the zero-second delay condition in Buffalo et al., 1998a. Model performance is 100% accurate for all experiments, as can be seen in Supplemental Figure S1 (top row) as well as Fig. 2 at x=1. We note that pixel-level representations also achieve 100% accuracy on all non-diagnostic experiments, an independent validation of our model-based exclusion criterion.

**Supplementary Figure S3:**
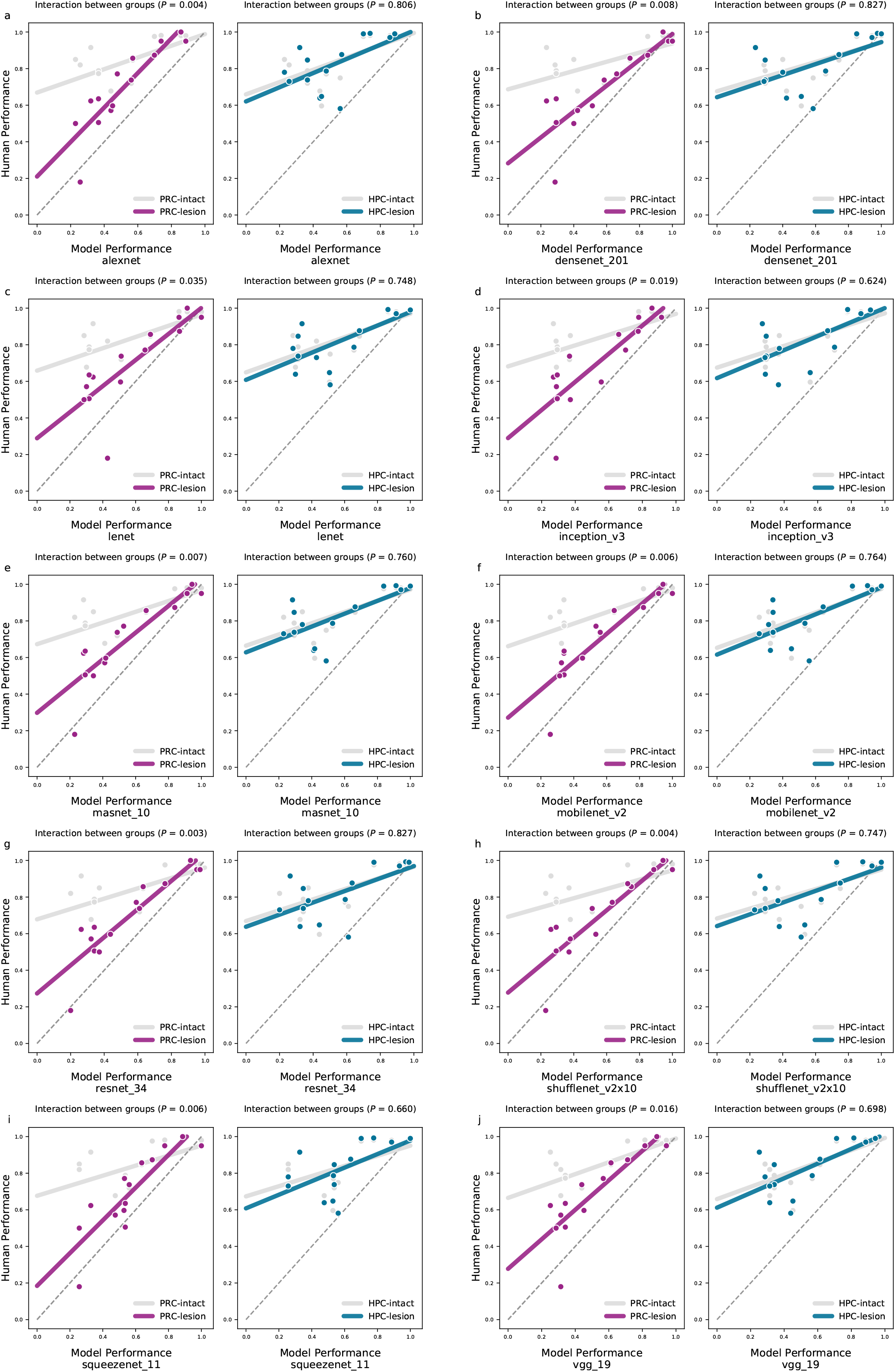
Retrospective results are consistent across all models evaluated, Related to Figure 2. Each pair of images recapitulates the analysis performed in Fig. 2, using a different instance of this convolutional model class, e.g. alexnet (a) and densenet 201 (b). As in Fig. 2, we situate PRC (left in subplot) and hippocampal (right in subplot) groups in relation to model performance. Across all architectures we observe a significant interaction between PRC-lesion and -intact performance, and the absence of a significant interaction between hippocampal-lesion and -intact performance. These interactions are described for each group, for each model, above each plot. Numerous models (e.g. ‘squeezenet’) have a better quantitative fit to PRC-lesioned performance than the model reported in the main analysis. We caution against interpreting these differences, given the low experimental *N* s within the retrospective analysis. We simply note that the same statistical relationship observed in the main text is present across all models evaluated.

**Supplementary Figure S4:**
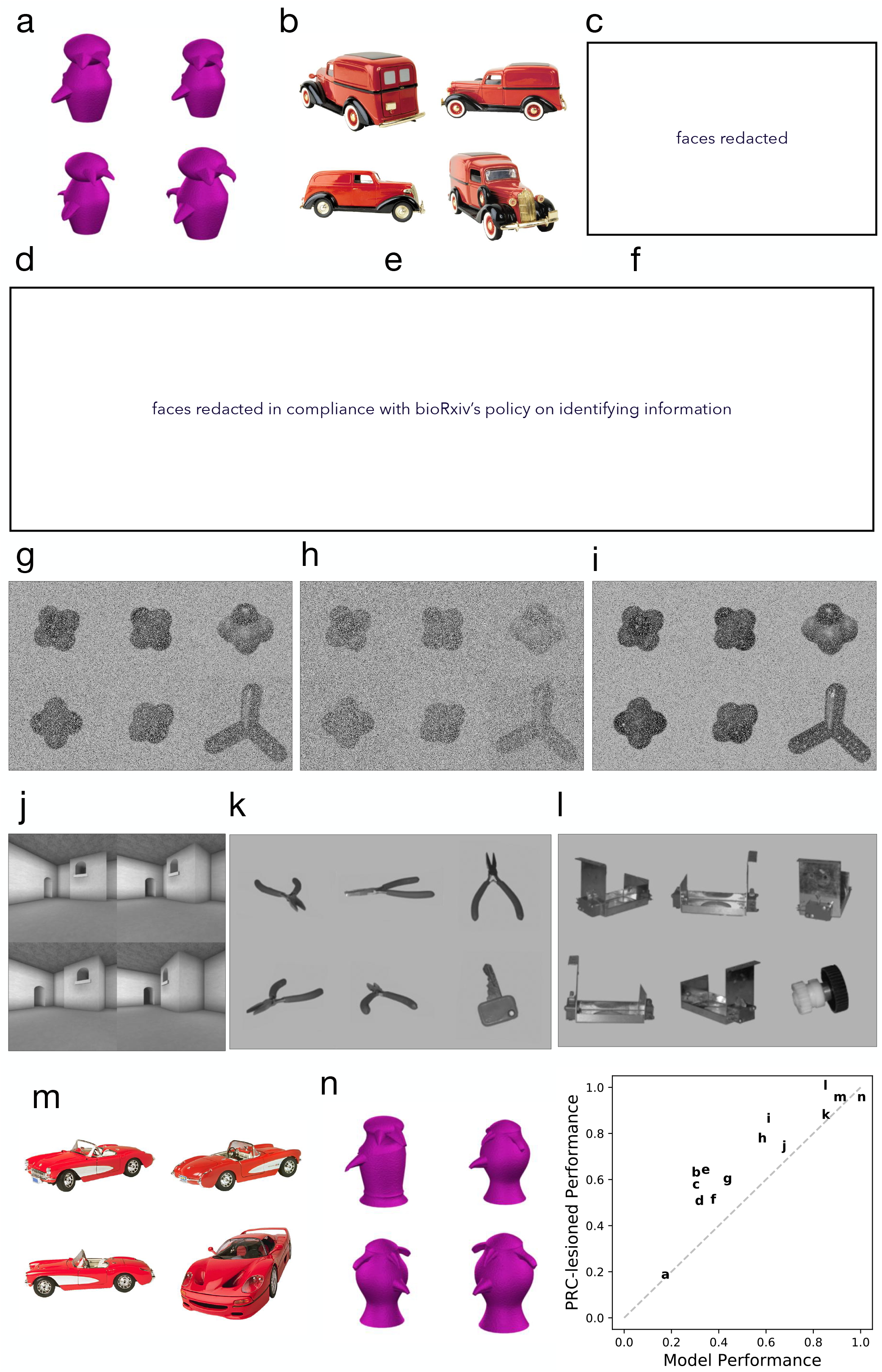
An example trial from each diagnostic experiment, Related to Figure 2. Experiments are in ascending order, in relation to model performance, with the lowest performing experiment at top left. **(a)** High Ambiguity novel objects in Barense et al., 2007, **(b)** High Ambiguity Familiar Objects in Barense et al., 2007, **(c)** Faces experiment one in Lee et al., 2005a, **(d)** Faces in Stark and Squire, 2000, **(e)** Faces experiment in Lee et al., 2005a, **(f)** Different View Faces in Lee et al., 2006, **(g)** Snow5 in Stark and Squire, 2000, **(h)** Snow6 in Stark and Squire, 2000, **(i)** Snow 4 in Stark and Squire, 2000, **(j)** Different View Scenes in Lee et al., 2005a, **(k)** Familiar objects in Lee et al., 2006, **(l)** Novel Objects in Lee et al., 2006, **(m)** Familiar Low Ambiguity Objects in Barense et al., 2007, **(n)** Novel Low Ambiguity Objects in Barense et al., 2007. The relationship between model and PRC-lesioned performance across experiments (a-n) is plotted in lower right.

**Supplementary Figure S5:**
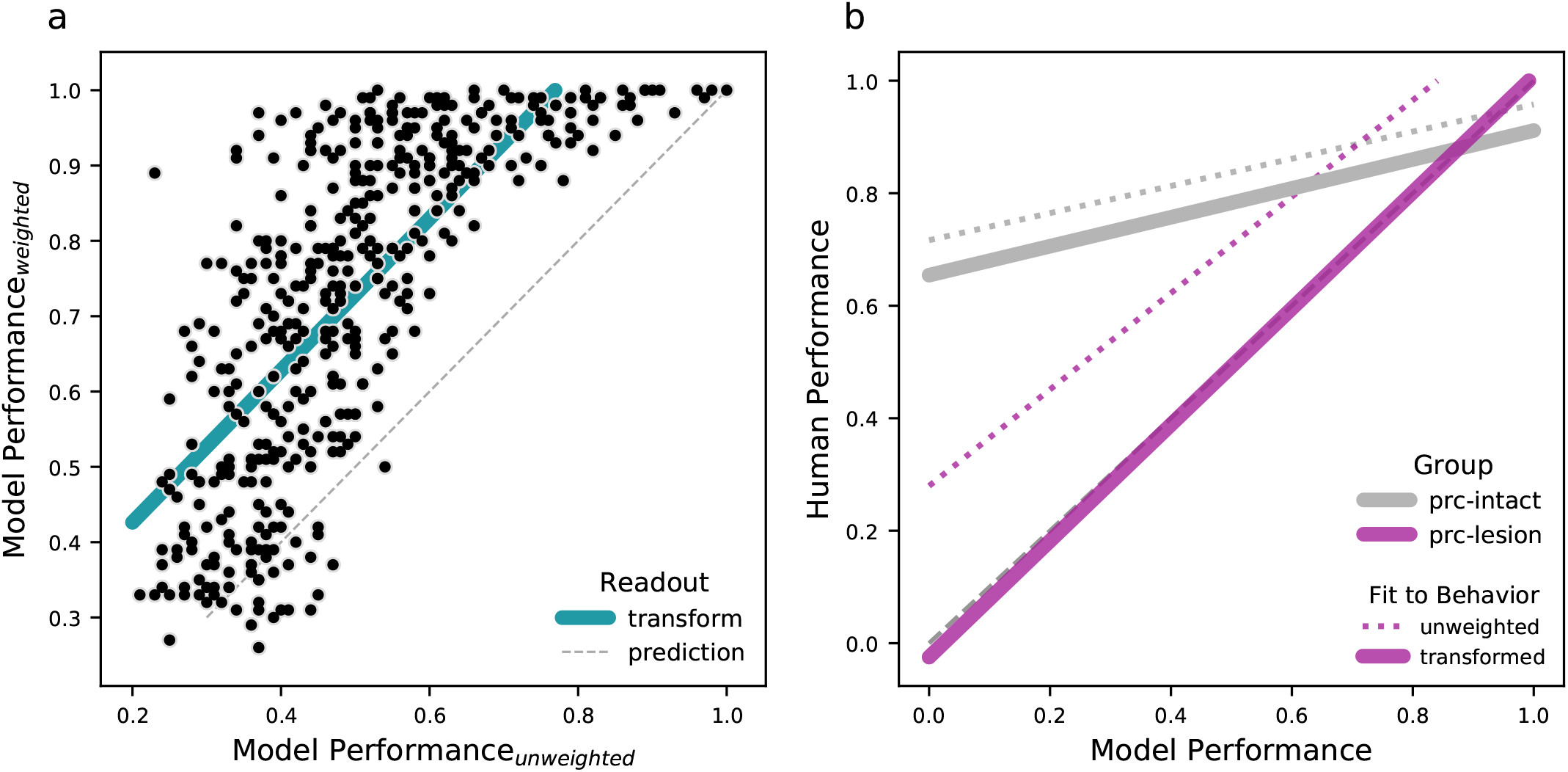
Correcting for readout type improves correspondence between model performance and PRC-lesioned behavior in retrospective analysis, Related to Figure 4 and 2. The interaction between PRC-lesion and -intact performance is central to our evaluation of the perceptual-mnemonic hypothesis. Nonetheless, there is a mean difference between PRC-lesioned and model performance, with PRC-lesioned participants performing significantly better (20%; Fig. 2a). We attribute this to four potential sources of variance. First, low sample size in the retrospective dataset encourages caution when interpreting these results. Second, these data are collected from human participants with naturally occurring PRC-lesions; incomplete PRC lesions may result in partially preserved PRC-relevant functions (Suzuki, 2009). Third, we observe modest variance in the absolute fit to PRC-lesioned accuracy across model architectures, with some model variants demonstrating an absolute performance level that approaches that of PRC-lesioned humans (Supplemental Fig. S3). Finally, underperformance may be a consequence of model readout. The uniform (i.e. distance-based) readout we employ in the retrospective analysis offers a stringent test of human-model correspondence, while a weighted, linear readout (which are standard practice in the literature) would be expected to better approximate human performance (Rajalingham et al., 2018). To evaluate this possibility, we use the novel stimulus set to compare **(a)** an unweighted linear (i.e. distance-based) readout of model responses (x axis), as per the original retrospective dataset analysis, alongside model performance supported by a weighted, linear readout of model responses, learned through a leave-one-out cross-validation strategy (y axis). As expected, the weighted, linear readout significantly outperforms the distance metric. We learn the transformation that projects the unweighted performance into the performance expected for the same stimuli using a learned, weighted readout (green). **(b)** Using the transform learned in (a), we project model performance supported by a uniform readout of model responses (i.e. the original retrospective analysis) into the accuracy that would be expected were it possible to learn a weighted readout on these stimuli. This improves the correspondence between model performance and human performance, suggesting that the mean difference between model performance and human performance may be due to the readout type. These results suggest that future efforts to better approximate PRC-lesioned performance must consider not only model instance, extent of lesion, and sample size, but also model readout as an important consideration with making human-model comparisons.

**Supplementary Figure S6:**
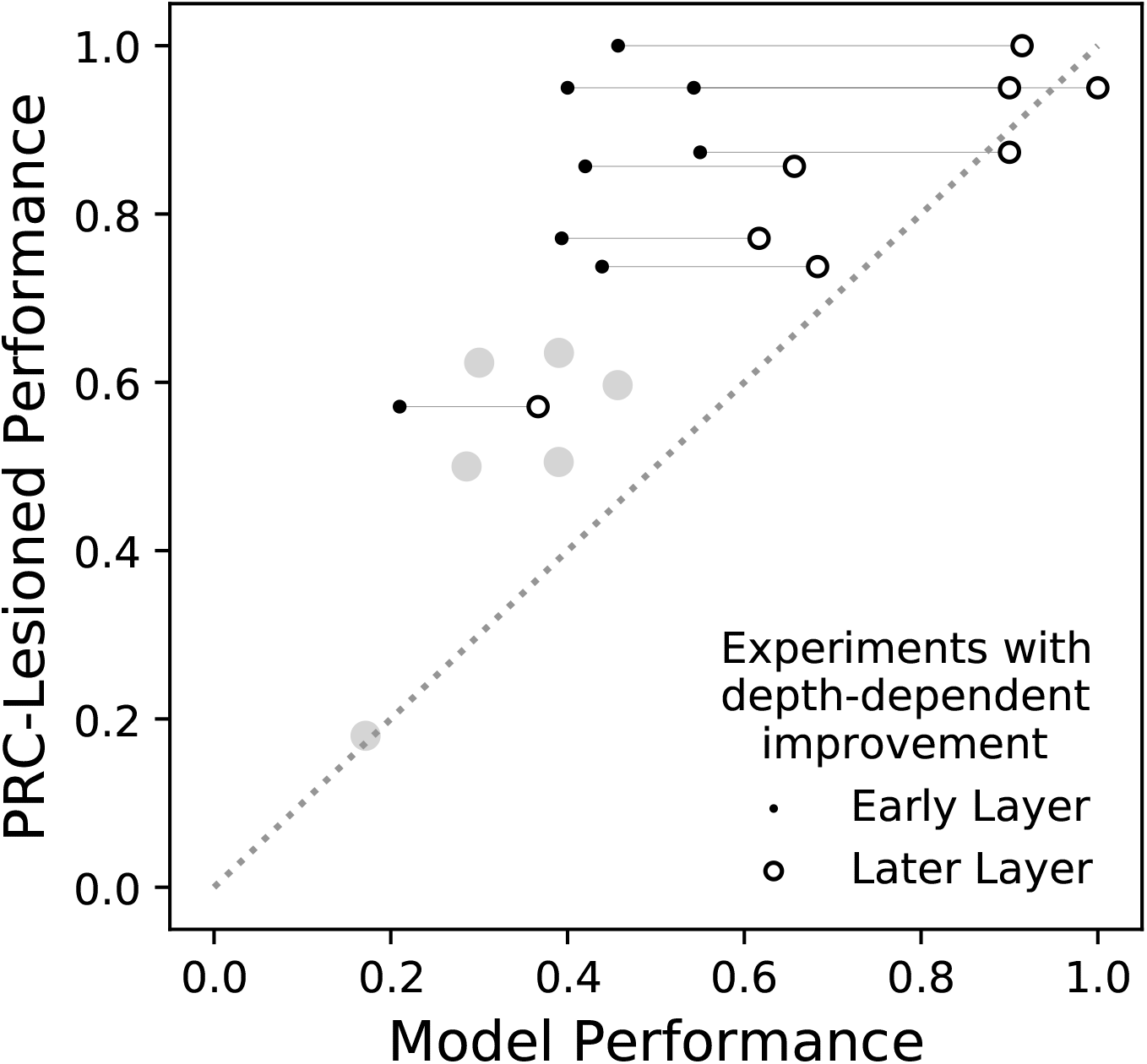
‘IT-like’ model layers fit PRC-lesioned performance, with depth-dependent improvements predominantly observed for ‘less complex’ stimuli, Related to Figure 2. Here we evaluate whether model performance on the retrospective dataset depends on the *readout layer* used to extract responses from the original model: Do deeper layers perform better? For each experiment, we determine whether there is a significant positive relationship between model performance and model depth. This is a complementary analysis to those performed in section 2.1.5: PRC-lesioned oddity performance appears to rely on high-level visual cortex. For each experiment in the retrospective analysis we evaluate model performance determined from each layer (not simply the most ‘IT-like’ model layer). We then compare these depth-dependent improvements with PRC-lesioned performance on each of these experiments (y axis). For those experiments on which there was a significant improvement with model depth (i.e. deeper/later model layers perform better then early model layers), on the × axis we visualize model performance from an early (e.g. V4-like, point) and later (e.g. IT-like, open circle) model layer, connected with a solid line. Those experiments where model performance does not improve with depth are visualized with a single grey point, corresponding to model performance from an IT-like layer. We find that model performance increases with depth for some experiments in the retrospective dataset (e.g. ‘Low Snow’ stimuli in Stark and Squire, 2000) but not others (e.g. ‘High Ambiguity Novel’ stimuli in Barense et al., 2007). We can separate experiments according to whether they exhibit depth-dependent improvements, then inspect the behavior of PRC-lesioned participants segmented along the same grouping. We find that PRC-lesioned participants perform significantly better on experiments that exhibited depth improvements (*µ* = .88) than on experiments that did not (*µ* = .52, *t*(6) = 5.17, *P* = .001). This latter group of experiments are those with the most substantial differences between PRC-lesioned and and -intact behaviors. Model performance increased for those experiments where PRC-lesioned subjects performed better. This quantitatively better fit to PRC-lesioned performance from more IT-like layers is suggestive that PRC-lesioned performance on the retrospective dataset is reliant on high-level visual cortex.

**Supplementary Figure S7:**
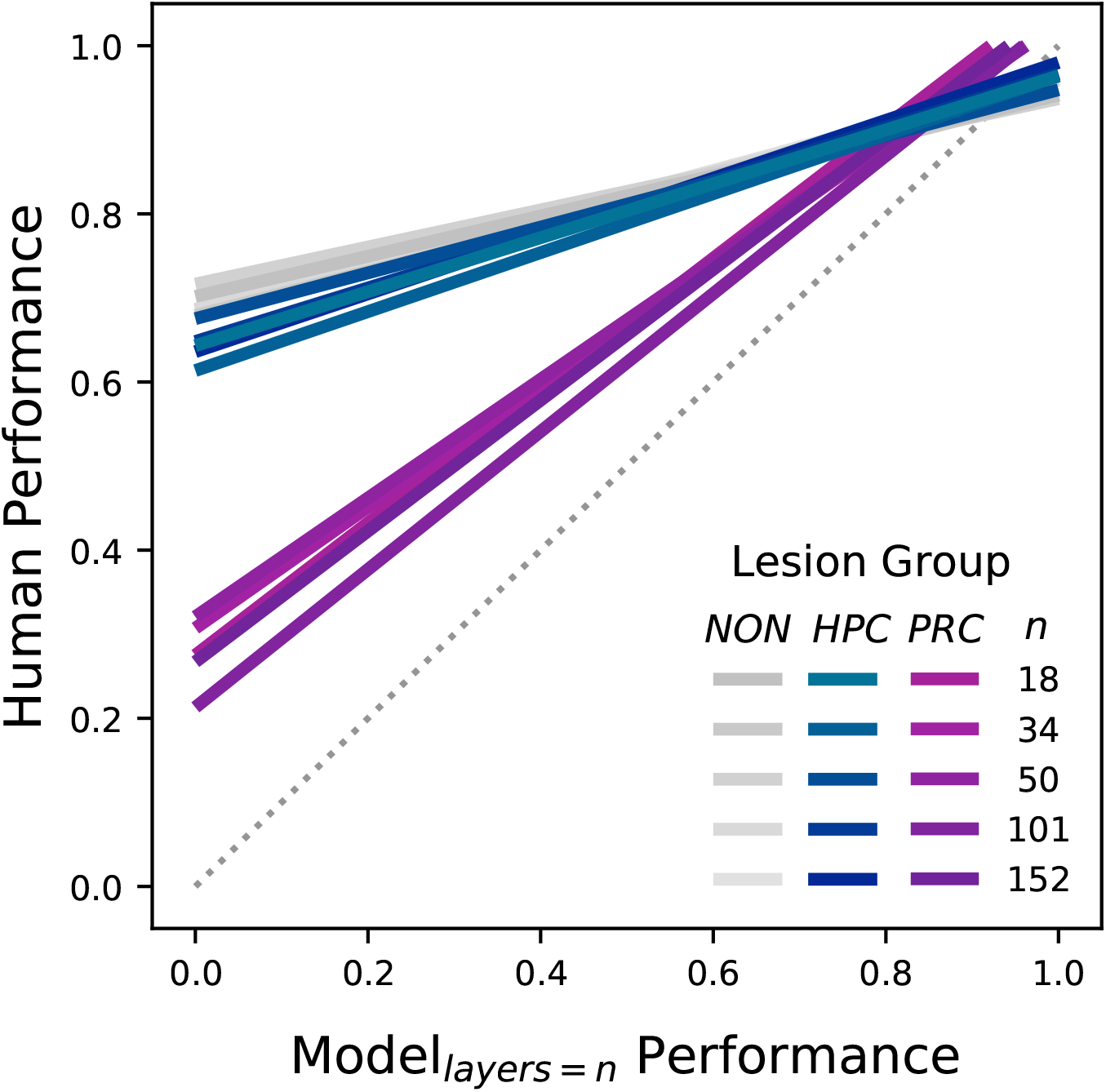
Increasing model depth does not improve performance on ‘more complex’ stimuli, or improve fit to PRC-intact behavior, Related to Figure 2. Our previous analyses (Supplemental Fig. S6) demonstrate that model performance improves when using stimulus responses extracted from deeper (i.e. later) model layers. Interestingly, this depth-dependent improvement was observed predominantly for experiments with ‘less complex’ stimuli. This leaves open the possibility that using models with more layers (i.e. increasing model depth) will lead to increased model performance on even the most ‘complex’ experimental stimuli. To evaluate this possibility, here we determine whether a simple modification to existing VVS-like models enables PRC-dependent behaviors: we systematically vary model architecture by increasing the number of layers (i.e. model depth). This approach has been shown to increase performance on computer vision benchmarks (He et al., 2016; Krizhevsky et al., 2012). We recruit a family of deep residual neural networks (i.e. ‘resnets’, He et al., 2016) optimized to perform object classification on a large-scale image classification task (ImageNet, Deng et al., 2009). This enables us to preserve the same computational motif across models while increasing the number of layers from 18 to 152. We implement this analysis using pretrained architectures from pytorch’s model zoo, and conduct the retrospective analysis (STAR Methods: Model Performance on Retrospective Dataset) using the penultimate layer as the readout used to determine model performance. We then compare model performance to PRC-intact (greys), HPC-lesioned (blues) and PRC-lesioned (purples) behavior for each model. Solid lines correspond to the best fit across all experiments. Across all resnets, our results are consistent with our original findings observed in Fig. 2. Critically, there is a significant interaction between PRC-intact and -lesioned participants in all models (e.g. 152 layers: *β* = .54, *F* (3, 24) = 3.97, *P* = 6×10^*-*4^). We find that adding layers does not improve performance on more ‘complex’ experiments, and deeper models do not perform these behaviors more like PRC-intact participants. These results corroborate findings from the lesion studies: Achieving PRC-dependent behaviors may require a computational motif distinct from these VVS-like architectures.

## Notes

### Competing Interest Statement

The authors have declared no competing interest.

## References

Aggleton, J. P., & Brown, M. W. (2006). Interleaving brain systems for episodic and recognition memory. Trends Cogn. Sci., 10, 455–463. 10.1016/j.tics.2006.08.003

Barense, M. D., Gaffan, D., & Graham, K. S. (2007). The human medial temporal lobe processes online representations of complex objects. Neuropsychologia, 45, 2963–2974. 10.1016/jzz.neuropsychologia.2007.05.023

Bashivan, P., Kar, K., & DiCarlo, J. J. (2019). Neural population control via deep image synthesis. Science, 364, eaav9436. 10.1126/science.aav9436

Brown, T. I., Staresina, B. P., & Wagner, A. D. (2015). Noninvasive functional and anatomical imaging of the human medial temporal lobe. Cold Spring Harb. Perspect. Biol., 7, a021840. 10.1101/cshperspect.a021840

Buckley, M. J., Booth, M. C., Rolls, E. T., & Gaffan, D. (2001). Selective perceptual impairments after perirhinal cortex ablation. J. Neurosci., 21, 9824–9836. 10.1523/JNEUROSCI.21-24-09824.2001

Buckley, M. J., & Gaffan, D. (1997). Impairment of visual object-discrimination learning after perirhinal cortex ablation. Behav. Neurosci., 111, 467. 10.1037//0735-7044.111.3.467

Buckley, M. J., & Gaffan, D. (1998a). Perirhinal cortex ablation impairs configural learning and paired–associate learning equally. Neuropsychologia, 36, 535–546. 10.1016/s0028-3932(97)00120-6

Buckley, M. J., & Gaffan, D. (1998b). Perirhinal cortex ablation impairs visual object identification. J. Neurosci., 18, 2268–2275. 10.1523/JNEUROSCI.18-06-02268.1998

Buffalo, E. A., Ramus, S. J., Clark, R. E., Teng, E., Squire, L. R., & Zola, S. M. (1999). Dissociation between the effects of damage to perirhinal cortex and area te. Learn. Mem., 6, 572–599. 10.1101/lm.6.6.572

Buffalo, E. A., Reber, P. J., & Squire, L. R. (1998a). The human perirhinal cortex and recognition memory. Hippocampus, 8, 330–339. 10.1002/(SICI)1098-1063(1998)8:4%3C330::AID-HIPO3%3E3.0.CO;2-L

Buffalo, E. A., Stefanacci, L., Squire, L. R., & Zola, S. M. (1998b). A reexamination of the concur-rent discrimination learning task: The importance of anterior inferotemporal cortex, area te. Behav. Neurosci., 112, 3. 10.1037//0735-7044.112.1.3

Bussey, T. J., Saksida, L. M., & Murray, E. A. (2003). Impairments in visual discrimination after perirhinal cortex lesions: Testing ‘declarative’ vs. ‘perceptual-mnemonic’ views of perirhinal cortex function. Eur. J. Neurosci., 17, 649–660. 10.1046/j.1460-9568.2003.02475.x

Bussey, T. J., Saksida, L. M., & Murray, E. A. (2002). Perirhinal cortex resolves feature ambiguity in complex visual discriminations. Eur. J. Neurosci., 15, 365–374. 10.1046/j.0953-816x.2001.01851.x

Bussey, T. J., Saksida, L. M., & Murray, E. A. (2005). The perceptual-mnemonic/feature conjunc-tion model of perirhinal cortex function. Q. J. Exp. Psychol. B., 58, 269–282. 10.1080/02724990544000004

De Leeuw, J. R. (2015). Jspsych: A javascript library for creating behavioral experiments in a web browser. Behav. Res. Methods., 47, 1–12. 10.3758/s13428-014-0458-y

Deng, J., Dong, W., Socher, R., Li, L.-J., Li, K., & Fei-Fei, L. (2009). Imagenet: A large-scale hierarchical image database. 2009 IEEE Conf. Comp. Vis. Patt. Rec., 248–255. 10.1109/CVPR.2009.5206848

DiCarlo, J. J., Zoccolan, D., & Rust, N. C. (2012). How does the brain solve visual object recogni-tion? Neuron, 73, 415–434. 10.1016/j.neuron.2012.01.010

Eacott, M., Gaffan, D., & Murray, E. A. (1994). Preserved recognition memory for small sets, and impaired stimulus identification for large sets, following rhinal cortex ablations in monkeys. Eur. J. Neurosci., 6, 1466–1478. 10.1111/j.1460-9568.1994.tb01008.x

Eichenbaum, H., & Cohen, N. J. (1993). Memory, amnesia, and the hippocampal system. MIT press. 10.1136/jnnp.58.1.128-a

Felleman, D. J., & Van, D. E. (1991). Distributed hierarchical processing in the primate cerebral cortex. Cereb. Cortex, 1, 1–47. 10.1093/cercor/1.1.1

Gaffan, D., & Murray, E. A. (1992). Monkeys (macaca fascicularis) with rhinal cortex ablations succeed in object discrimination learning despite 24-hr intertrial intervals and fail at match-ing to sample despite double sample presentations. Behav. Neurosci., 106, 30–38. 10.1037//0735-7044.106.1.30

He, K., Zhang, X., Ren, S., & Sun, J. (2016). Deep residual learning for image recognition. 2016 IEEE Conf. Comp. Vis. Patt. Rec., 770–778. 10.1109/CVPR.2016.90

Huang, G., Liu, Z., Van Der Maaten, L., & Weinberger, K. Q. (2017). Densely connected convolu-tional networks, 4700–4708. 10.1109/CVPR.2017.243

Iandola, F. N., Han, S., Moskewicz, M. W., Ashraf, K., Dally, W. J., & Keutzer, K. (2016). Squeezenet: Alexnet-level accuracy with 50x fewer parameters and¡ 0.5 mb model size. arXiv preprint.

Inhoff, M. C., Heusser, A. C., Tambini, A., Martin, C. B., O’Neil, E. B., K«ohler, S., Meager, M. R., Blackmon, K., Vazquez, B., Devinsky, O., et al. (2019). Understanding perirhinal contributions to perception and memory: Evidence through the lens of selective perirhinal damage. Neuropsychologia, 124, 9–18. 10.1016/j.neuropsychologia.2018.12.020

Jolly, E., & Chang, L. J. (2019). The flatland fallacy: Moving beyond low–dimensional thinking. Top. Cogn. Sci., 11, 433–454. 10.1111/tops.12404

Knutson, A. R., Hopkins, R. O., & Squire, L. R. (2012). Visual discrimination performance, memory, and medial temporal lobe function. Proc. Natl. Acad. Sci. USA., 109, 13106–13111. 10.1073/pnas.1208876109

Krizhevsky, A., Sutskever, I., & Hinton, G. E. (2012). Imagenet classification with deep convolu-tional neural networks. Adv. Neur. Inf. Pro. Sys., 25, 1097–1105. 10.1145/3065386

Lee, A. C., Buckley, M. J., Gaffan, D., Emery, T., Hodges, J. R., & Graham, K. S. (2006). Dif-ferentiating the roles of the hippocampus and perirhinal cortex in processes beyond longterm declarative memory: A double dissociation in dementia. J. Neurosci., 26, 5198–5203. 10.1523/JNEUROSCI.3157-05.2006

Lee, A. C., Buckley, M. J., Pegman, S. J., Spiers, H., Scahill, V. L., Gaffan, D., Bussey, T. J., Davies, R. R., Kapur, N., Hodges, J. R., et al. (2005a). Specialization in the medial temporal lobe for processing of objects and scenes. Hippocampus, 15, 782–797. 10.1002/hipo.20101

Lee, A. C., Bussey, T. J., Murray, E. A., Saksida, L. M., Epstein, R. A., Kapur, N., Hodges, J. R., & Graham, K. S. (2005b). Perceptual deficits in amnesia: Challenging the medial temporal lobe ‘mnemonic’ view. Neuropsychologia, 43, 1–11. 10.1016/j.neuropsychologia.2004.07.017

Levy, D. A., Shrager, Y., & Squire, L. R. (2005). Intact visual discrimination of complex and feature-ambiguous stimuli in the absence of perirhinal cortex. Learn. Mem., 12, 61–66. 10.1101/lm.84405

Majaj, N. J., Hong, H., Solomon, E. A., & DiCarlo, J. J. (2015). Simple learned weighted sums of inferior temporal neuronal firing rates accurately predict human core object recognition performance. J. Neurosci., 35, 13402–13418. 10.1523/JNEUROSCI.5181-14.2015

Meunier, M., Bachevalier, J., Mishkin, M., & Murray, E. (1993). Effects on visual recognition of combined and separate ablations of the entorhinal and perirhinal cortex in rhesus monkeys. J. Neurosci., 13, 5418–5432. 10.1523/JNEUROSCI.13-12-05418.1993

Miyashita, Y. (2019). Perirhinal circuits for memory processing. Nat. Rev. Neurosci., 20, 577–592. 10.1038/s41583-019-0213-6

Murray, E. A., & Baxter, M. G. (2006). Cognitive neuroscience and nonhuman primates: Lesion studies. MIT Press.

Murray, E. A., & Bussey, T. J. (1999). Perceptual–mnemonic functions of the perirhinal cortex. Trends. Cogn. Sci., 3, 142–151. 10.1016/s1364-6613(99)01303-0

Murray, E. A., Bussey, T. J., & Saksida, L. M. (2007). Visual perception and memory: A new view of medial temporal lobe function in primates and rodents. Annu. Rev. Neurosci., 30, 99–122. 10.1146/annurev.neuro.29.051605.113046

Murray, E. A., & Wise, S. P. (2012). Why is there a special issue on perirhinal cortex in a journal called hippocampus? the perirhinal cortex in historical perspective. Hippocampus, 22, 1941–1951. 10.1002/hipo.22055

Paszke, A., Gross, S., Chintala, S., Chanan, G., Yang, E., DeVito, Z., Lin, Z., Desmaison, A., Antiga, L., & Lerer, A. (2017). Automatic differentiation in pytorch. “

Rajalingham, R., Issa, E. B., Bashivan, P., Kar, K., Schmidt, K., & DiCarlo, J. J. (2018). Largescale, high-resolution comparison of the core visual object recognition behavior of humans, monkeys, and state-of-the-art deep artificial neural networks. J. Neurosci., 38, 7255–7269.10.1523/JNEUROSCI.0388-18.2018

Schrimpf, M., Kubilius, J., Lee, M. J., Murty, N. A. R., Ajemian, R., & DiCarlo, J. J. (2020). Integrative benchmarking to advance neurally mechanistic models of human intelligence. Neuron. 10.1016/j.neuron.2020.07.040

Scoville, W. B., & Milner, B. (1957). Loss of recent memory after bilateral hippocampal lesions. J. Neurol. Neurosurg. Psychiatry, 20, 11. 10.1136/jnnp.20.1.11

Shrager, Y., Gold, J. J., Hopkins, R. O., & Squire, L. R. (2006). Intact visual perception in memory-impaired patients with medial temporal lobe lesions. Journal of Neuroscience, 26, 2235–2240. 10.1523/JNEUROSCI.4792-05.2006

Simonyan, K., & Zisserman, A. (2015). Very deep convolutional networks for large-scale image recognition. ICLR 2015.

Squire, L. R., Shrager, Y., & Levy, D. A. (2006). Lack of evidence for a role of medial temporal lobe structures in visual perception. Learn. Mem., 13, 106–107. 10.1101/lm.178406

Squire, L. R., Stark, C. E., & Clark, R. E. (2004). The medial temporal lobe. Annu. Rev. Neurosci., 27, 279–306. 10.1146/annurev.neuro.27.070203.144130

Stark, C. E., & Squire, L. R. (2000). Intact visual perceptual discrimination in humans in the absence of perirhinal cortex. Learn. Mem., 7, 273–278. 10.1101/lm.35000

Suzuki, W. A. (2009). Perception and the medial temporal lobe: Evaluating the current evidence. Neuron, 61, 657–666. 10.1016/j.neuron.2009.02.008

Suzuki, W. A., & Baxter, M. G. (2009). Memory, perception, and the medial temporal lobe: A synthesis of opinions. Neuron, 61, 678–679. 10.1016/j.neuron.2009.02.009

Ullman, S. et al. (1996). High-level vision: Object recognition and visual cognition (Vol. 2). MIT press Cambridge, MA.

Yamins, D. L., Hong, H., Cadieu, C. F., Solomon, E. A., Seibert, D., & DiCarlo, J. J. (2014). Performance-optimized hierarchical models predict neural responses in higher visual cortex. Proc. Natl. Acad. Sci. USA., 111, 8619–8624. 10.1073/pnas.1403112111

